# Adaptive Charge Modulation Enables Focal, Selective Spinal Cord Stimulation

**DOI:** 10.64898/2026.06.16.732691

**Authors:** Ritwik Vatsyayan, Fadi Khoury, Tara S. Porter, Eleni Sinopoulou, Hadi F. Dayeh, Samantha M. Russman, Rhea Montgomery-Walsh, Kaushik Shukla, Helen Saad, Andrew M. Bourhis, Alec Salcido, Carly Shevinsky, Jihwan Lee, Karen J. Tonsfeldt, Akira Nagamori, Hoi Sang U, David M. Roth, Drew A. Hall, Sharona Ben-Haim, Eiman Azim, Tony Yaksh, Alexander Khalessi, Mark Tuszynski, Shadi A. Dayeh

**Author notes:** Contributing Authors.

## Abstract

Clinical neuromodulation primarily employs near-field low frequency electrical stimulation to activate neurons in the immediate vicinity of the electrode. We introduce Adaptive Charge Modulation (ACM), a spatiotemporal, charge-balanced stimulation strategy that focuses activation at deep tissue sites distant from the stimulating contacts. We apply ACM to stimulate deep regions of the spinal cord, distant from dorsal root entry zones which are preferentially activated during low frequency stimulation (LFS). ACM applies multipolar, biphasic rectangular pulses to exceed activation thresholds in deeper neuronal populations, with reduced surface activation, potentially due to high-frequency suppression of neural activity. In epidural spinal cord stimulation in rats, ACM achieved single-muscle selectivity among fourteen monitored muscles with minimal co-activation of other muscles. Using simultaneous, high spatiotemporal resolution, 2,112-channel brain-spine recordings, we characterized the response latencies and pathways consistent with focal recruitment at depth. We observed chronic stability of the electrode and ACM in freely behaving animals over 68 days post-implantation. By enabling focal activation with epidural surface electrodes, ACM may expand the reach and precision of neuromodulation and neural interfaces.

## Introduction

Electrical stimulation of the central nervous system has transformed clinical care by enabling targeted modulation of dysfunctional circuits in both the brain and the spinal cord. Clinically validated modalities span deep-brain stimulation for movement disorders^1,2^, Parkinson’s disease^3,4^, depression^5,6^, obsessive-compulsive disorder^7,8^ and chronic pain^9,10^; epidural spinal cord stimulation for pain^11^; closed-loop responsive neurostimulation for focal epilepsy^12,13^ and for volitional motor recovery after paralysis^1,14^.

Recent reviews and trials have motivated the development of novel stimulation waveforms including patterned, burst, high-frequency, and optimized shapes to improve depth selectivity, efficacy, and energy efficiency relative to conventional tonic stimulation^15–18^.

Focal activation is especially difficult in epidural low frequency stimulation (LFS) of the spinal cord, where the conductive cerebrospinal fluid (CSF) layer shunts current and biases recruitment toward large-diameter dorsal root fibers near the dorsal root entry zone rather than deeper segmental pools^19,20^. To improve spatial selectivity, prior research has explored conformal, high-density thin-film epidural arrays that enable finer multipolar steering^21^, as well as more invasive intraspinal microstimulation (ISMS) to engage spinal interneuronal circuits directly^22^. In the brain, “temporal interference” stimulation uses offset high-frequency sinusoids to generate a low frequency envelope at depth – an elegant, non-invasive attempt to create focal subsurface hot-spots^23^. Yet across these strategies, rehabilitative-grade focality remains limited: targeted LFS paradigms that restore movement in primates and humans primarily achieve function by selectively engaging dorsal roots and, thus, their downstream synaptic targets rather than producing tightly localized activation in deeper spinal pools^24^.

In this study, we present a new stimulation paradigm, Adaptive Charge Modulation (ACM), to selectively stimulate neuronal pools at depth within the spinal cord using surface epidural contacts. ACM delivers two high-frequency, biphasic rectangular pulse trains in a bipolar configuration from contacts flanking a desired subsurface locus; parameters (frequency, phase width and amplitude) are adaptively tuned so that charge per phase remain subthreshold at the neighborhood of the stimulation contacts (at the spinal surface), yet exceeds the depolarization/activation threshold at the focal point in the depth of the spinal cord. The square wave interference yields selective, more naturalistic stimulation mechanisms with sharp transient potential changes at the rising and falling edges of the square waves, enabling rapid polarization and depolarization of the neuronal membrane, consistent with physiological activation^25^.

We performed mechanistic sweeps over the parameter space to define the operating regime that produces focal activation in the rat spinal cord. Selectivity was quantified by comparing response amplitudes across fourteen lower-limb muscles (seven per side) using a recruitment-selectivity metric. To probe pathways and parameter limits, we recorded muscle activity via electromyography (EMG) and cortical responses via electrocorticography (ECoG) during ACM delivery. To mitigate CSF shunting and lateral current spread, we implemented a concentric-ring, multipolar electrode geometry that constrains the epidural potential and produces a narrow, axially directed field lobe toward the intended target. Finite-element modeling in COMSOL coupled with NEURON simulations predicted ACM-driven focusing and steerability across spinal segments^26,27^. We validated motor-pool engagement using c-FOS (cellular FBJ osteosarcoma oncogene) and NeuN (neuronal nuclei, stained with RBFOX3) immunohistochemistry in spinal cord sections, contrasting ACM with conventional low frequency stimulation^28,29^. Finally, we demonstrated chronic applicability by evoking right forepaw movements in awake, freely moving rats 9 days post-implantation.

## Results

### ACM evokes focal responses

We placed platinum nanorod (PtNR) spinal electrodes with both stimulating and recording contacts at the T13-L1 vertebral junction on the spinal cord^21^. The electrode array had 64 recording contacts of diameter D=30 μm with an impedance at 1 kHz of 24.4 kΩ ± 15.3 kΩ (mean ± standard deviation). The stimulating contacts had a diameter D=100 μm with 1 kHz impedance of 11.1 kΩ ± 5.5 kΩ^30^. The stimulating contacts had 30 μm wide concentric ground rings around them-these served as the return contact for stimulation^31^ (Fig. 1(A)). We used biphasic stimulating currents in the range of 50 - 200 μA, previously established to be within the Charge Injection Capacity (CIC) for PtNR contacts^32^, pulse widths of 100, 300 and 500 μs, and frequencies of 968.1 and 1000 Hz, and recorded EMG activity bilaterally in lower limb muscles using a pair of needle electrodes inserted into each target muscle. We monitored a total of 14 lower limb muscles, 7 on each side of the body, to capture a wide range of activation.

**Figure 1.**
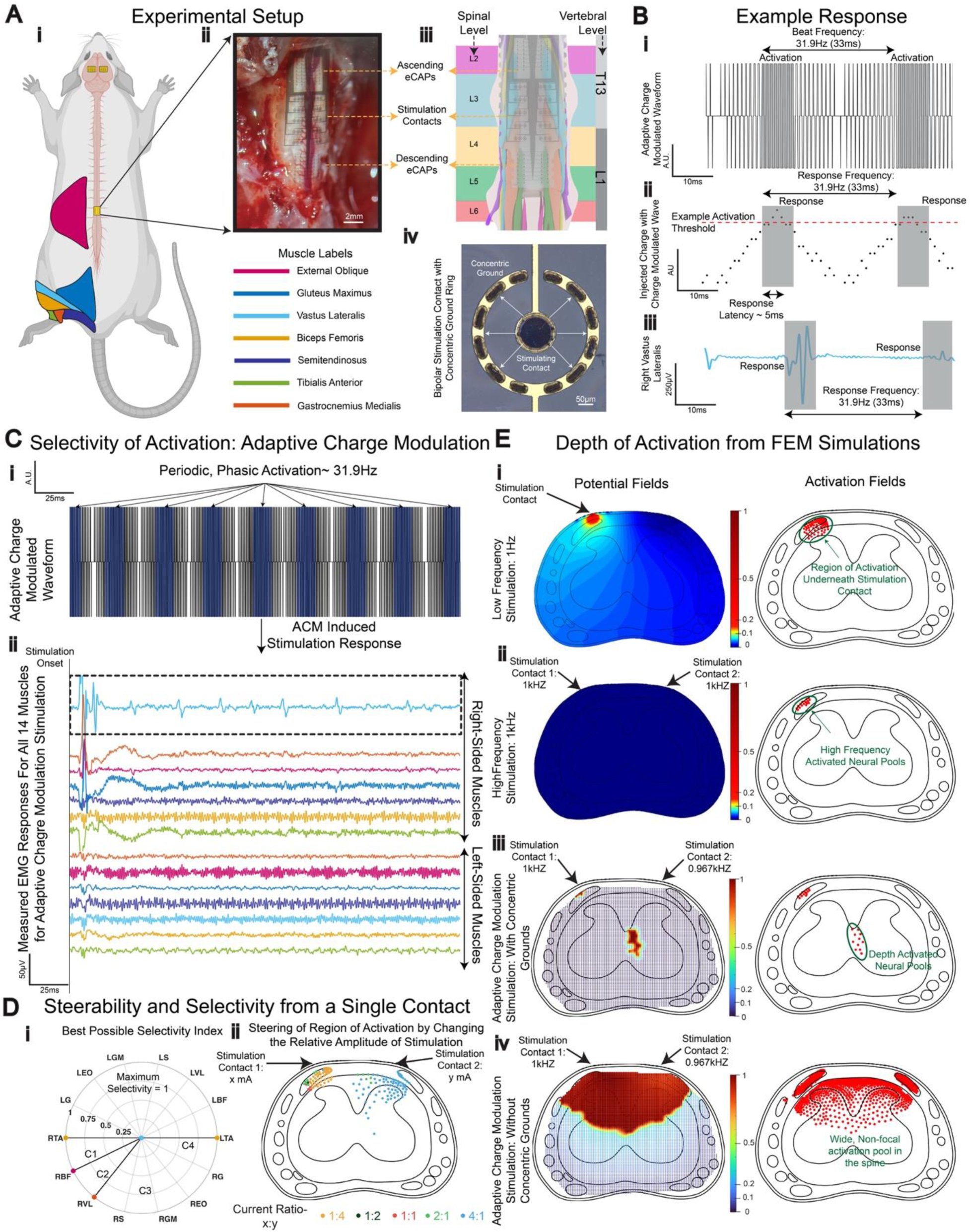
Experimental setup for ACM. (A) The experimental setup for spinal stimulation in the rat, highlighting the monitored muscles on the left-side (the same right-side muscles were also monitored). The inset shows an actual electrode placed conformally on the rat’s spinal cord. Aligned with this experimental placement is an illustration of the contact distributions across the L2-L5 spinal levels on a 3-dimensional model built from reconstructed spinal MRI images^21^. A single stimulation contact with the corresponding concentric ground is shown in the bottom-right image, captured under an optical microscope. (B) The resultant wave is generated from interference, with the corresponding magnitude of the charge injected in a single phase of the wave. The bottom row shows an example response waveform to stimulation for a random stimulation condition, with the response periodically repeating at the beat frequency. (C) A full 250 ms sequence of ACM waveform, with the elicited EMG response measured across all muscles, showing a strong first response to stimulation, and subsequent selective interference-targeted response to stimulation. (D) On the left, the best selectivity index achievable with a pair of stimulation contacts is obtained by modulating the relative amplitudes and pulse widths of stimulation. Four unique combinations of stimulation conditions were implemented, represented by C1-C4, to generate these responses. The corresponding EMG plots are shown in Supplementary Figure 2, along with the applied stimulation conditions required to produce the activation. The right panel shows the physical range of steerability achievable by varying the relative stimulation amplitude, as predicted by COMSOL simulations. (E) FEM simulations showing the activation region for LFS, high frequency stimulation, ACM and ACM without concentric ground, with the corresponding neuron activations calculated using NEURON, illustrating that activation at a sub-surface locus is best achieved with ACM. The color bar for the FEM simulations shows the electric potential in tissue, normalized to the maximum amplitude observed in tissue.

For a given set of biphasic pulses with a frequency offset, constructive interference in subsurface regions gives rise to pulse trains with a beat frequency equal to the offset frequency (Fig. 1(B)). As a result, the injected charge with ACM will periodically exceed activation thresholds at the beat frequency, yielding muscle responses locked to it (Fig. 1(B)). Under specific stimulation conditions applied to only 4 contact pairs, we selectively activate 7 (3 on the left side and 4 on the right side) out of the 14 monitored muscles (Supplementary Table 1). To illustrate this selectivity, we show that only the right Vastus Lateralis (RVL) exhibited activation among 14 monitored muscles (Fig. 1C). Stimulation with a pair of contacts (100 μA/ 500 μs/1000 Hz and150 μA/ 500 μs/968.1 Hz) generated a beat frequency of approximately 32 Hz, resulting in periodic activation. We defined EMG activation as responses with a signal amplitude exceeding either 5 standard deviations above the muscle’s baseline mean, or 25 μV for noisier recordings, whichever threshold was lower, as detailed in the Methods section. EMG activity below the activation threshold is weakly present in some muscles (e.g., Left Tibialis Anterior, (Fig. 1(C))).

To quantify the focality of ACM stimulation, we measured selectivity in muscle activation by calculating the relative amplitude of a given muscle’s response compared to the response of all muscles in response to stimulation. This metric is defined as the Selectivity Index (SI) of the muscle and is adapted from a previously used metric for muscle recruitment selectivity in response to spinal cord stimulation^33^. The first stimulation response, known as a transient/phasic response, arises from direct activation of muscle fibers rather than by ACM and is absent for subsequent stimulation pulses. We verified this by comparing responses to the same stimulation currents and pulse widths, both with and without an offset in the stimulation frequency. We observed that the first response in both conditions was almost identical, with no statistically significant differences (Supplementary Table 1). The subsequent activations seen in ACM were not observed with the no-offset stimulation (Supplementary Table 2). For a given set of stimulation parameters on the same pair of contacts, we achieved a SI=1 representing focal recruitment for the Right Tibialis Anterior (RTA), Right Biceps Femoris (RBF), Right Vastus Lateralis (RVL), and Left Tibialis Anterior (LTA) (Fig. 1(D), panel i, with the corresponding EMG plots shown in Supplementary Figure 1). We reproduced the selective stimulation of individual muscles in n=3 rats (Supplementary Figure 2).

Targeting focal responses to stimulation has long been a major consideration in the field of LFS in the spinal cord^33–38^. With ACM, the neural pools in the vicinity of the stimulation contact experience high amplitude, high frequency stimulation, with minimal modulation from the second stimulation wave. Therefore, these neural pools do not respond to stimulation^39^. Due to volumetric conduction within the spinal cord, stimulation pulses from two contacts constructively interfere in the depth of the spinal cord, and only the neural pools in the region of optimal interference respond to stimulation, creating a focal response region. Since the EMG and cortical activations that we typically record do not rule out activation of neural pools in the vicinity of the stimulation contact, we performed finite element model (FEM) simulations using COMSOL Multiphysics to model the spread of the electric field in the spinal cord. We also modeled the response of Ia afferent neurons in the spinal cord using NEURON simulation package to predict activation thresholds to stimulation^26^. The simulation setup is shown in Supplementary Figure 3 (COMSOL setup) and Supplementary Figure 4 (NEURON setup). The simulations show that for the typical surface stimulation, the electric field is concentrated in the vicinity of the stimulation contact, and we see activation largely in the surface neuron pools underneath the stimulation contact. With ACM, the neurons in the vicinity of the high-frequency stimulation contact only get activated once, to the initial stimulation burst, as shown in Supplementary Figure 4(C). This initial single response is identical to the response we observe when we stimulate both the contacts at 1 kHz, both with simulations, and in actual experiments (Fig. 1(E) and Supplementary Figure 5).

### Concentric Grounds Help Focus the Subsurface Fields

LFS generally lacks focality in the recruiting neural pools in the depth of the spinal cord, due to the spread of current in the CSF beneath the dura and due to activation in the surface-conducting layers of the spinal cord^26,40–42^. Typically, a bipolar setup involves the flow of current between two symmetric, circular contacts placed on the surface of the spinal cord. The current spread can be mitigated by placing a current return contact (ground) as close as possible to the stimulation contact, limiting the activated neural tissue to the surface layers between.

To achieve focality, we designed our stimulation electrode comprising a stimulating contact and an adjacent concentric ground ring serving as the bipolar current return contact, as shown in Fig. 1(A), panel iv. In this setup, the lateral spread of current is limited, and the injected current travels spatially as an elliptical wavefront shaped by the adjacent ground in both lateral directions along the circumference, medio-lateral axis and along the rostro-caudal axis of the spinal cord. We modeled the effect of the concentric ground through a set of FEM simulations using COMSOL Multiphysics and modeled the potential gradient in the spinal cord^1,43^. With concentric grounds (Fig. 1(E), panel iii), the stimulation is focal within the spinal cord tissue, whereas for the same stimulation conditions and without concentric ground (Fig. 1(E), panel iv), the full surface of the spinal cord is stimulated. These simulation results demonstrate that concentric grounds effectively confine the electric fields generated during stimulation, an essential feature of ACM.

### ACM with pulse width and amplitude control

The ACM for subsurface stimulation provides three main parameters for selective activation: the frequency, the amplitude, and the width of the pulse for the two interfering square waves. By tuning the pulse width alone, we have shown spatially constructive interference resulting in supra-threshold charge activation and corresponding evoked activity (Fig. 2(A)). With a 500 μs pulse width (close to 50% duty cycle), the charge modulated wave has a complete interference profile and an almost complete temporal overlap, with no amplitude intermodulation. However, as we reduce the duty cycle (300 and 100 μs pulse widths), we introduce an additional amplitude modulated component to the target waveform within the constructive interference regime. These subtle changes impact the integrated charge density and the transition between stimulating and non-stimulating conditions.

**Figure 2.**
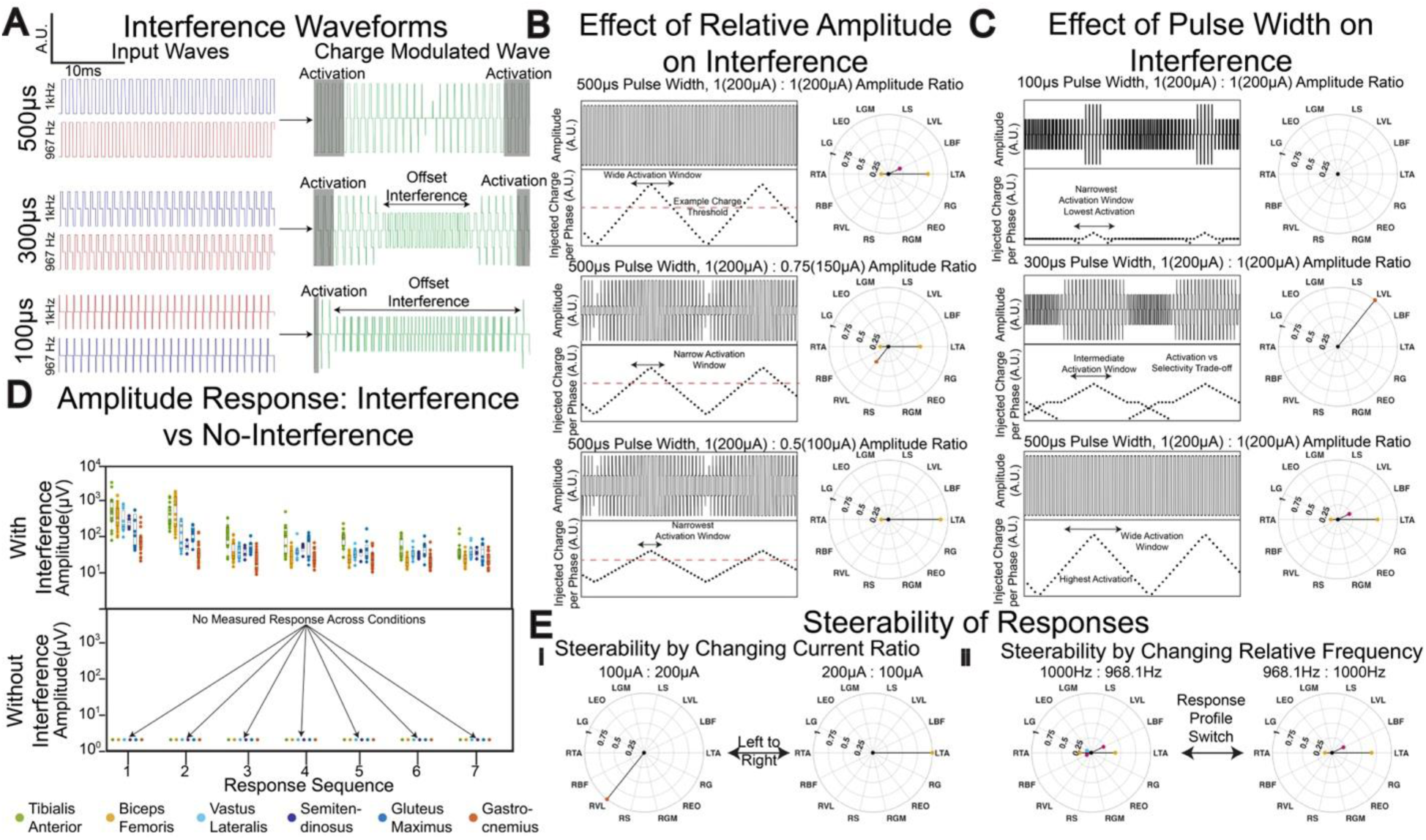
Principle of activation of ACM. (A) Example interference waveforms created from the stimulation waveforms from the two contacts, for 3 different pulse widths of the high-frequency waves:100, 300 and 500 μs. (B) The impact of the relative amplitude of stimulation on the interference waveform and the corresponding injected charge per beat cycle. The SI plot shows the experimental response profile for the corresponding conditions. (C) The impact of stimulation pulse width on the interference waveform, showing the more complex interference patterns generated, along with their corresponding experimental SI plots. (D) EMG response amplitudes for ACM and control (non-ACM; no frequency offset between injected pulses) stimulation conditions, with every ACM condition replicated with no frequency offset. (E) The EMG response amplitude comparing ACM and low frequency (LFS) stimulation, capturing the variation of subsequent response amplitudes to stimulation. This is a representative comparison; due to differences in setup, a direct comparison is not possible.

For the same pulse width and frequency, but with different stimulation amplitudes, unique waveforms with a critical charge exceeding the activation threshold can be obtained (Fig. 2(B), Supplementary Figure 6(A) shows the corresponding EMG responses). The combination of different pulse widths and amplitudes can also yield interference patterns that transition from activating conditions (larger pulse widths) to non-activating conditions (smaller pulse widths), as illustrated in Fig. 2(C) (Supplementary Figure 6(B) shows the corresponding EMG responses). Higher amplitudes typically create broader activation volumes as more charge is injected into tissue. By contrast, smaller amplitudes typically create narrower activation volumes. Similarly, wider pulse widths inject significantly more charge, creating a wider activation region. In contrast, the narrower pulse widths inject significantly lower charge and create a narrower window of activation. This leads to a trade-off between selectivity and activation, as shown in Fig. 2(C), panels ii and iii. The pulse width, therefore, serves as another tunable knob that can affect both the selectivity and the steerability of activation.

### Validating ACM Responses and Steerability and Selectivity of Activation

Electrical stimulation of excitable tissue is widely known to result in a nonlinearly proportional neural recruitment^44–49^. Thus, to effectively recreate multiple patterns of activation, a wide range of parameters, both in stimulation and electrode design may be required. To explore the achievable selectivity range for each contact pair, we varied each of the three critical parameters as discussed above. We varied the pulse widths between either 100, 300 or 500 μs. To limit the parameter space, both the interfering waves were set to the same pulse width. The relative amplitudes of stimulation at the two contacts were varied from 1:0.25 to 1:1 in steps of 0.25. For each contact pair, 96 unique stimulation parameter combinations were studied to explore the stimulation capabilities of ACM. Additionally, every stimulation parameter combination setting was also tested with a control, no offset stimulation, where the frequency of the injected wave from both pairs of contacts was set to 1 kHz, so that no interference would take place (Fig. 2(D), and the effect of offset vs no-offset stimulation is shown in Supplementary Figure 5).

Moreover, changing the stimulation parameters from the same pair of contacts allows us to move the window of activation in the spinal cord. Fig. 2(E), panel i shows the change in activation from the left side of the body, LTA, to the right side of the body, RVL, by interchanging the stimulation amplitude from the two contacts (150 μA: 100 μA to 100 μA: 150 μA). Similarly, changing the relative stimulation frequency of the two contacts changes the resultant waveform within the spinal cord, which in turn changes the window of activation, and the corresponding EMG response profile, as shown in Fig. 2(E), panel ii. In addition to selectivity, steerability of the region of response is therefore another strength of the proposed approach, with the same pair of contacts allowing a broader response profile.

The wide range of tested parameters enabled us to observe multiple response patterns in the target muscles (Fig. 3(B)). The wide parameter space afforded by ACM allows fine-tuned steering of activation fields in the spinal cord, and opens the door for the recreation of more naturalistic motion patterns in future studies. We tested ACM across 3 experiments on different animals with marginally different electrode placements and were able to achieve high selectivity and steerability in all 3 cases (Supplementary Figure 1). The electrode placement on the animal for these 3 cases, as well as all previous cases used for data collection, is shown in Supplementary Figure 7.

**Figure 3.**
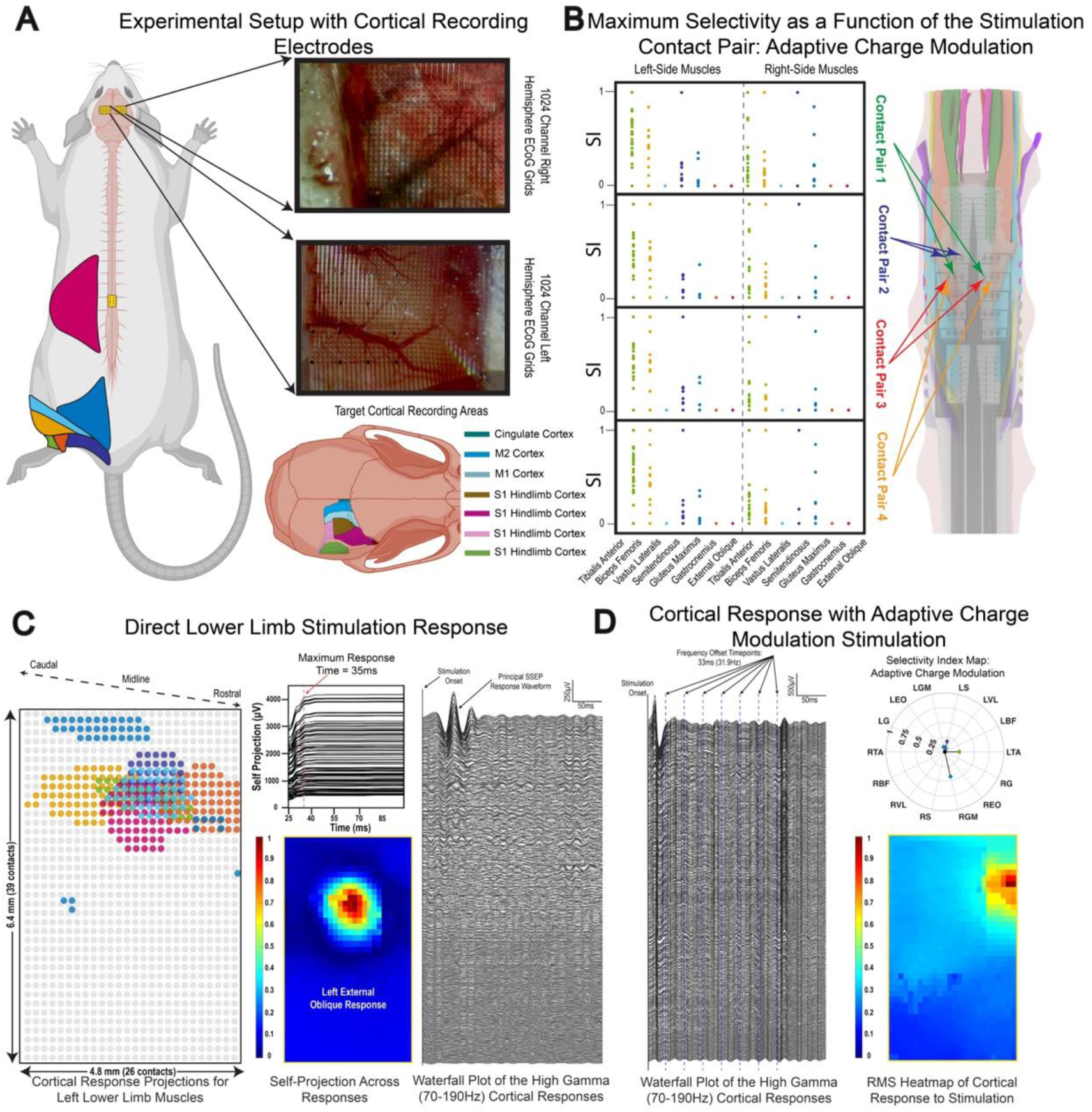
Parameter space of activation and cortical validation. (A) The experimental setup invoking cortical electrodes during ACM. Two 1024-channel ECoG arrays were placed relative to the midline, targeting the S1-M1 cortex in both hemispheres. (B) Selectivity Index response for each muscle for all stimulation conditions tested across a single experiment in ACM, showing the ability to achieve multiple levels of activation for muscles with a limited subset of stimulation contacts. (C) The validation of the placement of the cortical electrodes with evoked cortical activity from direct stimulation of the lower limb muscles. The response time on the cortex was calculated from the self-projection of responses, and the response region for each muscle was determined by thresholding these projections. The spatial heatmap is calculated from the z-scored RMS of the response to stimulating a single muscle (LEO). (D) Waterfall plot of cortical responses to ACM stimulation on a 1024-channel ECoG grid with the representative SI for the stimulation condition; the spatial heatmap is calculated with a z-scored RMS of the response.

To validate the response profiles from stimulation, we simultaneously recorded cortical and muscle responses elicited from spinal cord stimulation using ACM. We placed a 1024-channel thin film ECoG electrode array on each cortical hemisphere, approximately centered over the sensory-motor cortex^50^ (Fig. 3(A)). To verify electrode placement and validate responses from the target lower limb muscles, we first performed direct electrical stimulation of the seven left-sided muscles through the bipolar EMG needle electrodes used for subsequent recording. Fig. 3(C) shows the measured neural responses from direct electrical stimulation for each muscle. Using the high-density ECoG array enables us to localize cortical response areas for each muscle with very high precision. Next, we measured the cortical response profile for the ACM stimulation. Since we cannot directly measure neural activation in the spinal cord due to stimulation artifacts, we measured the cortical response to stimulation as a second response endpoint. Fig. 3(D) shows the cortical response profile for ACM. Notably, we registered cortical response epochs for each interference response time-window.

### The Minimum Frequency for ACM using Parametric LFS Stimulation and Simultaneous Brain, Spinal cord, and Muscle Recordings

Previous studies have shown that high-frequency stimulation inhibits neural responses, but the mechanisms and pathways underlying this inhibition remain unclear. We performed an array of mechanistic studies to evaluate spinal cord stimulation efficacy as a function of stimulation design parameters, recording simultaneously from the brain, muscles, and the spinal cord surface.

Fig.4(A) shows the ECoG response waveforms in the brain, measured in the sensory and motor cortices^51^ (Supplementary Figure 8 shows the placement of the cortical grids in the last 6 experiments of the study). The average duration of the cortical response to direct monopolar LFS, with a Pt needle electrode placed along the spinal midline rostral to the stimulation contact acting as the return contact, is approximately 25 ms. This implies an upper limit of ∼50 Hz for the maximum detectable stimulation-evoked response frequency. At 2 Hz, repeated responses to LFS are observed more consistently in the sensory cortex, while the response timing shows some offset in the motor cortex. As the frequency of the LFS increases, the responses and their timing become less consistent. However, as the response amplitude decreases, the spatial extent of the response becomes more focal (Fig, 4(D)). At 50 Hz, the response to stimulation becomes much less consistent, with overlapping responses from consecutive stimulations almost completely hiding the response waveform (Fig. 4(A)). Fig. 4(E) shows the response cutoff frequency, averaged across the 10 most responsive cortical channels, indicating approximate cutoff frequencies of 70 and 75 Hz for motor and sensory responses, respectively.

**Figure 4.**
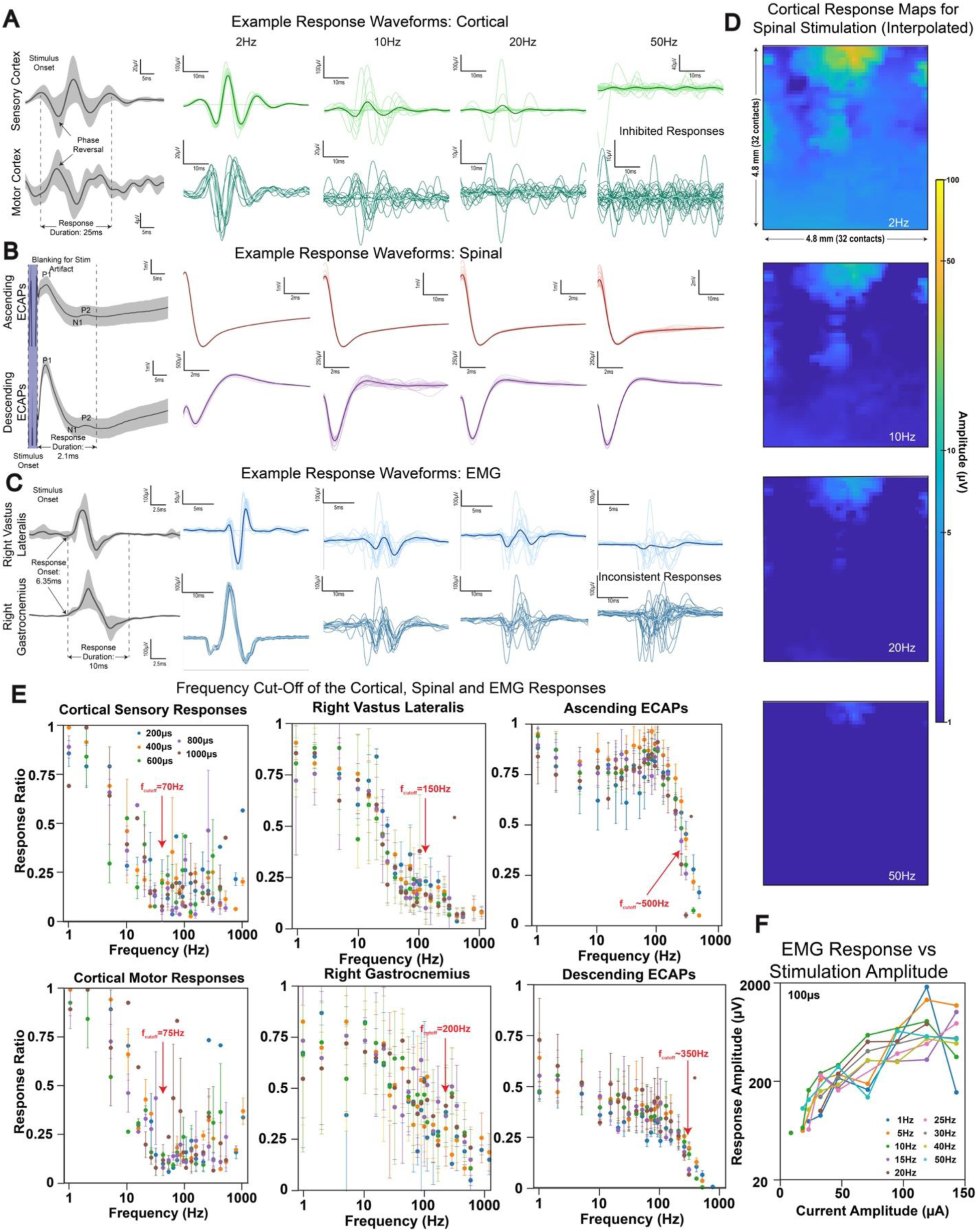
Mechanistic studies of the frequency response of LFS to determine focal ACM stimulation conditions. Example response waveforms, for the activity measured in the (A) cortex, (B) spinal cord and (C) muscles. The first panel shows the example response at 1Hz, with the response standard deviation of the responses shown with the grayed background. The other columns show the changes in the response waveform as a function of frequency. (D) The cortical response heatmap shows decreases in response amplitudes across the brain as stimulation frequency increases. The plots include a linear interpolation of data between functional channels to account for broken channels. (E) The frequency cutoff for each target variable, showing the eventual suppression of responses at high-frequency. (F) The observed variation in the EMG response, as a function of the stimulation amplitude.

Fig. 4(B) shows the responses recorded from the rostro-caudal surfaces of the spinal cord for LFS. The ascending evoked Compound Action Potentials (ECAPs), representative of sensory signals, are measured from electrode contacts rostral to the stimulation contacts, whereas the descending ECAPs, representative of motor signals, are measured from electrode contacts caudal to the stimulation site. The spinal responses to stimulation, the first point of activation, are more consistent in both amplitude and timing, and have a significantly higher cutoff frequency than the cortical responses, closer to 500 Hz for ascending ECAPs. The presence of stimulation artifact limits the analysis of frequencies higher than 500 Hz, which is why spinal responses were not analyzed for ACM stimulation.

Fig. 4(C) shows the EMG response to LFS for two right-sided muscles. Similar to the cortical responses, the EMG responses are a few synapses down the primary activation pathway, so their response cutoff frequencies are lower than those observed in the spinal cord. The normalized response ratio falls below 0.25 close to 150 Hz for the Right Vastus Lateralis, and 200 Hz for the Right Gastrocnemius. Therefore, the stimulation frequency for the high-frequency square waves must be higher than the response cutoff frequency for each target vector along the neural pathway. This is primarily determined by the spinal response frequency, which is close to the response cutoff frequency of individual neurons. To avoid direct activation of neurons in the spinal cord and to enable focal sub-surface stimulation, our ACM stimulation frequencies used in this study were above 967 Hz.

Increasing the stimulation amplitude causes the response amplitude to increase, until the response amplitude plateaus around 2 mV, as shown in Fig. 4(F). Surface stimulation is widely understood to elicit responses from the nerve roots beneath the stimulation contact, and the resulting response profile therefore reflects direct activation of these nerve roots. The minimum and maximum stimulation currents used for ACM are within the tested windows. The EMG response amplitudes elicited through ACM, except for the first response, were consistently lower-even at higher stimulation amplitudes. One possible explanation is that these responses arise from activation of neuron pools located deeper within the spinal cord, where the electric field amplitudes are lower (Supplementary Figure 9). We observed that LFS with voltage-clamped sinusoidal pulses (Supplementary Figure 10(A)) generally tends to have a larger cutoff frequency of activation (Supplementary Figure 10(B)), though the exact mechanism is not well-understood.

### Extension and Stability of ACM in the Chronic Freely Moving Animal

We first validated the chronic stability of the ACM electrodes – necessary for clinical translation^52–54^ – by implanting 4 rats with cervical spinal implants target the upper limbs as illustrated in Fig. 5(A). The implanted electrode contacts were 100 μm in diameter with a starting acute *in vivo* impedance of ∼ 5 kΩ at 1kHz, which increased immediately post-implantation due to the local response of tissue around the implant site^55,56^, and then remained relatively stable and below 100 kΩ for all tested animals (Fig. 5(D)). The longest tested implant for a stable electrode was 68 days. We performed ACM on one freely moving rat post-implant targeting paw contraction (Fig. 5(E) and Supplementary Video 1) using fixed stimulation conditions (biphasic square pulse, pulse width 0.1ms, pulse period 1ms, continuous trains of 1 second, amplitude range from 0.5 to 0.6V based on overt movement threshold) and without running a parametric sweep to hit optimal upper limb targets.

**Figure 5.**
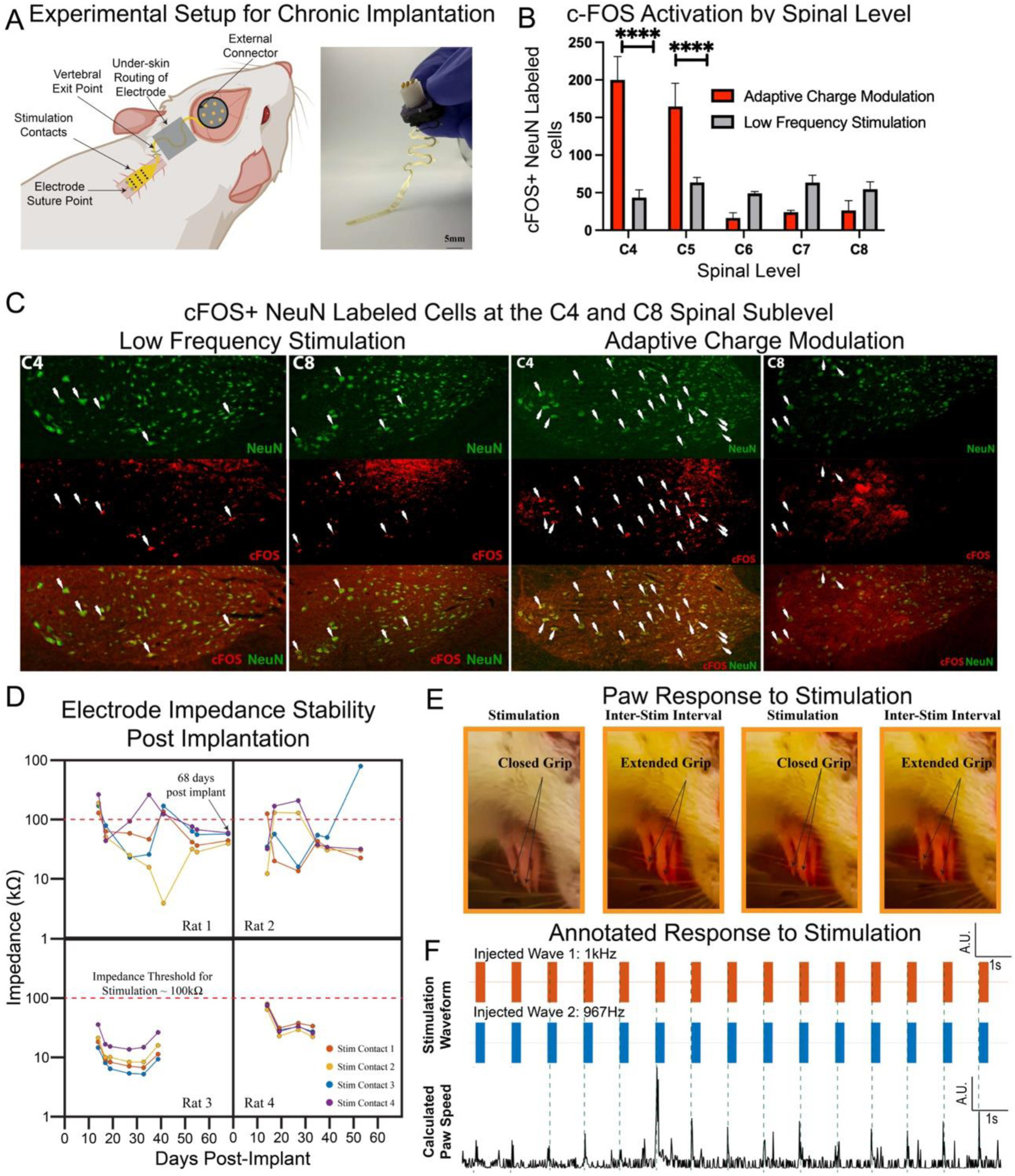
Validation of Sub-surface Stimulation with ACM and Extension of ACM to the Chronic Freely Moving Animal. (A) Experimental setup for the chronic experiments, showing the design of the chronic implantation system, with the inset showing the electrode design for the chronic experiment. (B) The variation in measured c-FOS+, NeuN labeled neurons at spinal levels C4-C8, for ACM and LFS. (**** is p < 0.0001, two-way ANOVA test) (C) The immunohistology images showing the c-FOS+, NeuN labeled neurons corresponding to (B), showing the variation between ACM and LFS. (D) The variation of impedance of the chronically implanted electrode over time, showing the stability of the electrode. (E) The measured motion of the paw in response to stimulation in a freely moving rat. The muscle contractions were confirmed using video analysis. (F) The response pattern was calculated by employing paw tracking, showing the motion of the paw in response to each stimulation pulse.

We repeated ACM at 1s intervals to cause a paw contraction, and the motion was captured using video. The electrode was allowed to stabilize for one-week post-implantation. The animal was then stimulated daily from days 6-9 post-implantation. Paw position was tracked using Adobe After Effects, and the movement speed was calculated from the tracked points for the behavior recorded on Day 9. Fig. 5(F) shows that the sharp increases in the speed of the paw coincided with the delivered stimulation, validating the functionality of ACM in the chronic freely moving animal. The longest implanted animal, Rat 1, was stimulated with the same ACM protocol up to 68 days post-implantation, with the electrode impedance remaining stable throughout the course of the experiment.

### Immunohistological Validation of ACM Subsurface Stimulation

To validate the extent and specificity of activation achievable with ACM stimulation, we performed terminal electrophysiology experiments and immunohistochemical staining on the stimulated spinal cords using c-FOS to identify the activated cells. We used 2 paradigms; ACM and LFS both paradigms focused on the upper spinal segments (C4 and C5). The electrode placement spanned from the C5 spinal level to C8 level of the cervical spine (Supplementary Figure 11(A) and (C)) and stimulation was intended to target the activation of the forelimb biceps and triceps. We performed both ACM and LFS for 30 minutes, with stimulation sequence repeated every 1s, in separate animals and prepared the tissue for staining (n = 1 for each experiment). The animals were euthanized immediately at the end of the experiment and subsequently perfused with 4% PFA (time to perfusion did not exceed 7 minutes for any experiment). The spinal tissue was then processed and sectioned. Sections from the C4 to the C8 spinal levels were stained with c-FOS and NeuN.

To identify activated neurons in the spinal cord, we analyzed the count of c-FOS+ NeuN labeled neurons, as shown in Fig. 5(B). We observe a significantly higher activation at the C4 and C5 spinal levels with ACM (two-way ANOVA; F(4,19)=52.73, P<0.0001). By contrast, LFS shows a wider distribution in activated c-FOS+ neurons from C4-C8. This indicates a greater focality in neuron activation with ACM at the desired level when compared against LFS, consistent with the improved selectivity observed with lower-limb recruitment.

## Discussion

### Mechanism of Activation of ACM

ACM works on the principle of spatiotemporal interference between two high frequency waves in the depth of the spinal cord. We demonstrated three tunable parameters for generating an interference pattern: the amplitude of stimulation from the two contacts, the pulse width of the underlying square waves and the frequency offset between the two waves. The parameter space is therefore large, and for this study, we have focused on exploring a limited subset for demonstrative purposes.

High frequency waves travel through a region of inactivation around the stimulation contact, typically at the entry of the nerve roots into the spinal cord^39,57^. This was confirmed by mechanistic frequency studies, as shown in Fig. 4(E), which demonstrate that the neural responses to stimulation of the spinal cord, and consequently the activity further down synaptically at the brain and the muscles, are suppressed at higher frequencies. At the region of interference with suprathreshold charge injection, we were able to create a region of activation deep within the spinal cord, without activating tissue directly adjacent to the stimulation contacts, effectively creating stimulation patterns that would otherwise require penetrating electrodes to the spinal tissue.

At the point of perfect interference, we create a repeating pattern of square waves with varying pulse widths, each injecting a fixed amount of charge per phase. We hypothesize that activation occurs in the spinal cord when the charge injected in a given phase exceeds the threshold for neuronal excitation, thereby eliciting a stimulation response that follows the spatiotemporal interference of the amplitude-modulated wave. This is best represented by the interference patterns shown in Fig. 2(A). The region of activation, however, is not limited to the area of perfect interference, which is why the same pair of contacts can simultaneously activate multiple muscles whose neuron pools lie in different regions in the depth of the spinal cord.

ACM allows us to control the interference pattern by modifying another important parameter in stimulation: the pulse width of the underlying waves. The higher pulse widths, closer to 50% duty cycle of the square wave, allow for a simpler, repeatable patterns of stimulation. The lower pulse widths create more interesting interference patterns since the duty cycle of the pulse is lower than 50%, which means one gets periods of both interference, and periods of complete offset in interference. The lower pulse width yields lower charge injected per phase which results in a smaller region of activation. However, with multiple temporal offsets in stimulation, the response frequency is less controllable – for applications that require consistent periodic responses – because it is possible to have multiple responsive waves as compared to higher pulse width stimulation.

One of the major advantages of this technique is the selectivity and the steerability that is achievable from the same pair of contacts on the spinal cord. The interference of two waves from two different contacts creates a more focal region of activation in the depth of the spinal cord, whereas reaching the depth of the spinal cord through LFS would create a much larger activation region, with limited focality. Across three different animal experiments, we were able to selectively stimulate a wide range of lower limb muscles from a very small combination of stimulation contacts.

Further, within the same pair of contacts, the region of activation can also be steered to multiple locations on both sides of the midline of the spinal cord, allowing us to stimulate a much wider range of muscles. By changing the injected current from the two stimulating contacts (2 independent parameters), the pulse width of stimulation (two independent parameters, although for this study, the pulse width from the two stimulating contacts was held equal), one can have a much wider range of activation patterns of lower limb muscles than would be possible with traditional surface stimulation. This could enable functional rehabilitation with much smaller electrodes, significantly improving surgical outcomes in spinal surgery and reducing the load on device and electrode design required to support stimulating electrodes.

In this work, we did not explore the role of frequency offset in the generating stimulation patterns. This was partially due to hardware limitations from our current stimulation system. The time step for a 30 kHz stimulation system is about 33 μs, which means that the closest frequency step we can get around our stimulation frequency of 1kHz is roughly 32 Hz. In the future, the development of clinical stimulation systems with higher time resolutions, amplitude control and channel count will enable finer frequency steps and more precise targeting within the spinal cord. This study was designed to robustly validate ACM for sub-surface focal and steerable stimulation using state-of-the art spinal stimulation and brain electrodes that are already part of an FDA-approved clinical trial, but additional future research is required for scale-up and to translate ACM for spinal cord therapies.

### The Principle of Spatiotemporal Interference

The principle of spatiotemporal interference to stimulate neurons in the depth of neural tissue, bypassing those neurons at the surface, has previously been demonstrated^23^, and is gaining increasing traction for potential application in non-invasive stimulation therapies^58–65^. This approach was first proposed for transcranial electrical stimulation, using sinusoidal electrical waves. The interference of sinusoidal waves to create an amplitude-modulated wave is one of the fundamental principles of long-distance communication systems and has previously been applied in neural stimulation to develop non-invasive stimulation systems.

In our approach, we used square waves rather than sinusoidal waves, driven primarily by our goal of recreating more naturalistic activation patterns in neurons^65^. For decades, neurostimulation has been driven using square waves, with the sharp transition edge enabling the opening and closing of ion channels in the neural membranes. Further, square waves allow exquisite control over the charge delivery and, consequently, over the spatiotemporal interference patterns. For example, traditional DBS therapies for brain network disorders such as epilepsy, Parkinson’s and depression, use square -pulse stimulation at frequencies as high as 200 Hz^66–69^, whereas in the spinal cord, high frequency stimulation has been used for pain therapies^70–72^.

In this work, we have integrated the design of a novel spatiotemporal interference paradigm with high-density, thin film electrodes, demonstrated to have long-term stability chronically in rodents. The high-density, low-impedance electrodes, coupled with the design flexibility offered by the stimulation paradigm, allow for a wide-parameter space to tune the activation in the spinal cord, and thereby extend the achievable recruitment patterns for stimulation, presenting a significant step forward in the design of stimulation therapies targeting neuron populations in the depth of the spinal cord and the brain.

### Limitations in the Current Study Design

While the current design allows us to explore the response space across a broad parameter space, we recognize that the proposed method has limitations that need to be resolved for potential clinical translation and use. The presence of the first response to high-frequency stimulation, which arises when neurons fire once before being inactivated by high-frequency stimulation, creates a potential side-effect that must be managed. While not covered in this study, we conjecture that this effect can be controlled by providing a ramped increase to the stimulation, allowing the generation of a high-frequency field at lower amplitudes that raises the threshold for activation of the neurons without directly activating the neurons, as has been shown previously with sinusoidal stimulation^23^. Alternatively, using lower pulse widths also controls the amount of charge initially injected, and can reduce the first activation amplitude.

This study was also limited to a single frequency of activation for spatiotemporal interference; the variation in response with stimulation frequency needs to be studied in greater detail. Further, while the stimulation setup is chosen to closely mimic available clinical systems, there can be general variability in the choice of stimulation system, including the time latency of the square wave’s rising edge and the temporal resolution of stimulation, which can create variability in responses. Additionally, the safety of high-frequency stimulation from a neural tissue perspective has not been extensively studied and can limit the amplitude and pulse width.

We attempted to measure the EMG responses from a broad range of muscles in the rat’s lower body, extending from the trunk to the lower body, all the way down to the gluteus muscles. This, however, is not representative of all motor neuron pools innervated in the spinal cord in the lumbo-thoracic region, and therefore, we cannot rule out other co-activated neural pools that might limit focal selectivity in practical applications. We also are unable to directly measure the activation of the dorsal nerve roots, and the exact time within the temporal interference window where the injected charge exceeds the activation threshold cannot be precisely determined. This means that secondary estimates of dorsal vs ventral activation in the spinal cord through latency calculation between stimulation and EMG responses cannot be used. Lastly, the validation of the long-term stability of the thin film electrode was performed in the cervical spinal cord to allow the electrode to be stabilized to the vertebral bones, and the connector to be placed on the skull. The development of a translational system for the lower spinal cord would require additional design components, not studied here.

## Conclusions

In this work, we presented a novel method to stimulate neural pools deep within the spinal cord, termed as adaptive charge modulation (ACM). We demonstrate high focality in the recruitment resolution achieved in anesthetized rats. We then performed mechanistic studies to determine the frequency thresholds for ACM and to explore how the frequency of stimulation impacts neural recruitment and how high-frequency interference shapes activation patterns. We then demonstrated the applicability of the method in a translatable, chronic setup by implanting and stimulating the spinal cord in an awake, freely moving animal. This technique has broad impact for a wide range of potential applications in neurological disorders of both the brain and the spinal cord and applicability towards future development of brain-computer interfaces.

## Acknowledgments

We acknowledge insightful discussions with Gert Cauwenberghs of UC San Diego. This work was performed, in part, at the San Diego Nanotechnology Infrastructure (SDNI) of UCSD, a member of the National Nanotechnology Coordinated Infrastructure, which is supported by the NSF (grant ECCS1542148). Tissue Technology Shared Resource is supported by a National Cancer Institute Cancer Center Support Grant (CCSG Grant P30CA23100).

## Author contributions

These authors contributed equally: Fadi Khoury and Tara S. Porter

Conceptualization: S.A.D., R.V., and S.M.R.

Method Development: R.V., F.K., T.S.P., E.S, H.F.D., S.M.R., R.M-W., K.S., H.S., A.M.B., J.L, K.J.T., A.N., H.S.U, D.R., D.A.H., S.B-H., E.A., T.Y., A.K., M.T., S.A.D.

Electrode Fabrication: R.V., F.K., T.S.P., K.S., S.M.R.

Experimental Data Collection: R.V., F.K., E.S., T.S.P., S.M.R., H.S.U., A.S., C.S.

Computational Simulations: R.V., F.K., H.F.D., R.M-W, H.S.

Investigation: R.V., F.K., T.S.P., E.S, H.F.D., H.S., H.S.U, D.R., M.T., S.A.D. Visualization: R.V., F.K., E.S, H.F.D, S.A.D.

Funding acquisition: E.S., S.B.-H, A.K., M.T., S.A.D. Project administration: S.A.D.

Supervision: S.A.D.

Writing – original draft: R.V., S.A.D.

Writing – review & editing: all co-authors.

## Data Material and Availability

All data are available in the main text or the supplementary materials, and will be uploaded to the DANDI Archive, per NIH funding mandates.

## Methods

### Stimulation Electrode Design and Fabrication for the Spinal Electrodes

The thin-film electrodes used in this study were fabricated at the UC San Diego Nanofabrication Facility (nano3). The polymer material used as the substrate for the acute experiments was Parylene-C, and for the chronic experiments was Polyimide. The electrode interface material used in all experiments is Platinum Nanorods (PtNR) that exhibits superior impedance and stability performance ideal for stimulation. The interface material was connected to the acquisition electronics using evaporated and encapsulated metal traces made of Chromium and Gold (20 nm/ 250 nm).

For acute experiments, the electrodes were heat-bonded to anisotropic-conductive film (ACF) tapes, which were connected to custom designed printed circuit boards (PCBs) that interfaced with the background system. The stimulation contact for the spinal electrodes had a diameter of 100μm. The concentric ground was designed with an inner radius of 170 μm and an outer radius of 200 μm. There were 6 rows of contacts, placed symmetrically either side of the mid-point of the electrode which was designed to go over the midline of the spinal cord. The impedance cutoff for the working contacts was considered from 5-30 kΩ at 1 kHz, measured post-placement against a stainless-steel ground contact placed in the animal’s back, approximately midline.

The recording contacts for the ECAPs in both the ascending and descending directions had a diameter of 30 μm and were arranged in a configuration of an 8 x 4 array (longitudinal x circumferential). Circumferentially, the contacts were spaced 1mm apart, and longitudinally, they were placed 500μm apart. The impedance cutoff for the recording contacts was considered from 10 – 100 kΩ at 1 kHz, measured against the same stainless steel ground contact used for stimulation, since both the recording and stimulation contacts shared the same acquisition system. The various iterations of the electrodes used, and the animal experiments they were used in, are shown in Supplementary Figure 12 and Supplementary Table 3, respectively.

For the chronic experiments, the electrodes were bonded directly to the custom designed PCBs using Silver epoxy (Ag epoxy). The electrode lead regions were designed using a wavy pattern that allowed the electrode leads to expand and contract as the electrode moved. This PCB was then housed in a 3-D printed structure made using biocompatible ABS polymer. This housing was then stabilized on the rat skull using dental cement. The impedance cutoff for a working stimulation contact in the chronic setup was considered from 10-100 kΩ at 1 kHz. The lower impedance limit was used to rule out contact breakdown or shorting between neighboring contacts, and the upper impedance limit was set to allow safe and below compliance injection of at least 50 μA (resulting in 5 V – near compliance – for a 100 kΩ impedance) at 100 μs pulse width.

### Recording Electrode Design and Fabrication for the Cortical and EMG Electrodes

We used ECoG electrodes for the cortical recording of responses in the rat brain. The cortical electrodes were also made using Platinum Nanorods (PtNR), and the substrate material used was Parylene-C. The electrode contact for recording had a diameter of 30 μm, and the inter-contact spacing was 160 μm (the electrode designs used are shown in Supplementary Figure 13). The thin-film electrodes were bonded using Ag epoxy to the acquisition electronics using an extender PCB. The extender PCB was connected to a custom-designed PCB that interfaces with the acquisition electronics.

For the muscle recordings, we use needle electrode pairs to record the EMG activity evoked from stimulation. The electrode material was stainless steel, and the twisted wire was connected to a custom-designed acquisition PCB through a touchproof connector.

### Data Acquisition Setup

Intan RHS stimulation controllers were used for stimulation across all experiments. The spinal electrodes used in the acute experiments were connected through custom PCBs to four Intan RHS2132 headstages. These headstages can simultaneously record and stimulate from 32 channels. The sampling frequency, which controls both the recording and stimulation timestep, was set to 30 kHz.

The cortical recording electrodes were connected to a custom acquisition PCB with embedded Intan RHD2164 high-density recording chips. An Intan 1024-channel RHD recording controller was used to simultaneously control 16 recording chips. The sampling frequency of the recording experiments was set to either 20 kHz or 30 kHz, depending on the length of the experiment.

The EMG recording electrodes were connected to an RHD2132 bipolar recording headstage that can record 16-bipolar recording pairs simultaneously.

The three systems had independent ground and reference electrodes, which allowed us to reduce the stimulation artifact picked up on the recording systems. All electrical systems, including the controllers, heart rate and anesthesia monitors, and the connected laptops, were run through noise-suppression systems to reduce the impact of 60 Hz line noise. All systems used in the experiment were time-locked using digital signals generated from the stimulation system. To run-through hundreds of stimulation parameters in an efficient manner, the stimulation controller was controlled using MATLAB, interfaced to the Intan RHX GUI.

### Data Analysis and Processing

#### EMG

The data acquired in the experiments was analyzed using MATLAB R2024a. For the EMG experiments, the signals were filtered between 10 and 500 Hz using a 2^nd^ order Butterworth bandpass filter^73^. To distinguish responses from noise, we defined a response as a signal with amplitude exceeding either 25 µV (≈10× the amplifier input-referred noise floor of 2.4 µV_RMS_) or 5 standard deviations above the mean baseline of a given channel, whichever threshold was lower. This combined criterion accounts for both the hardware noise floor and channel-specific variability, enabling sensitive yet robust detection across recordings. The maximum response duration for a single EMG response was 5 ms. The maximum peak-to-peak response detected across experiments was determined to be close to 2 mV.

Selectivity index for each muscle was calculated as a ratio of the response in the target muscle, vs the observed response in every other muscle, for that stimulation condition:

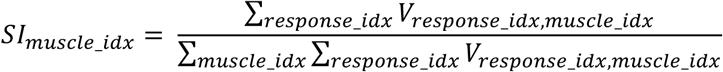

where, *V_response_number,muscle_index_* is the response calculated for a given muscle, and the response number can be greater than 1-for ACM stimulation we expect to find between 7-9 responses^33^. For ACM, the selectivity index was calculated by dropping the first response, which is typically the response generated from the direct activation of neural pools due to the first stimulation pulse.

For the comparison of the evoked response between frequency-offset (ACM) and no–frequency-offset stimulation, a two-sample t-test assuming unequal variances was performed. Significance thresholds were defined as follows: no significance (-) for *p* > 0.05; * for 0.05 ≥ *p* > 0.005; ** for 0.005 ≥ *p* > 0.0005; and *** for *p* ≤ 0.0005 (Supplementary Table 1 and Supplementary Table 2).

#### Cortical

The cortical data was filtered between 70 and 190 Hz using a 2^nd^ order Butterworth bandpass filter to capture the high-gamma neural activity. The common mode responses in the data were removed by performing Common Average Re-referencing.

#### Spinal

The spinal data, specifically to capture the ECAPs, was filtered between 10 and 500 Hz using a 2^nd^ order Butterworth bandpass filter^74,75^. The presence of a stimulation artifact post-stimulation made the analysis more challenging, and we did not specifically model out the stimulation artifact response.

Instead, the data chunk used in the analysis was considered from 1ms after stimulation, and the response waveform was re-averaged to have a zero-mean.

### FEM Simulations and NEURON Modeling

The FEM (finite element model) simulations of the spread of the electrical fields in the spinal cord were performed using COMSOL Multiphysics. An electric current (ec) model was used to calculate the electrodynamics of the electric fields in response to stimulation. The spinal cross-section used for simulation was generated from MRI cross-sections of the rat spinal cord, as published in a previous study from the lab. The same model was used to create the graphic of the spinal cord shown in Fig. 1(A).

The time-domain simulations from COMSOL were exported to a csv spreadsheet and imported into a python-script running the NEURON package. The simulations for activation from stimulation were performed assuming a spinal sensory Ia fiber is being simulated-in this study, we do not consider the spatial distribution of different neural pools. The model for the neuron used in simulation was adapted from previously published spinal cord stimulation neurons from the group of Marco Capogrosso.

### Vertebrate Animal Subjects and Surgical Procedures for Acute Studies

Adult male Sprague-Dawley rats (> 6 months old, weight range 750-1250 g, Charles Rivers Laboratories) were used for the acute parametric studies. General anesthesia was administered prior to surgery using 4% isoflurane, and the rats were fixed on a stereotaxic frame (Kopf Instruments). The anesthesia level was monitored using a rat paw pulse oximeter (Kent Scientific), and the isoflurane levels were reduced to 1.5-2.5% during surgery. A laminectomy was performed at the T13-L1 spinal levels, exposing the dorsal surface of the spinal cord without damaging the dura.

After surgery, the rat was transitioned to a ketamine/xylazine (90%/10%) intramuscular injection, re-dosed at every 20-30 mins. Body temperature was maintained between 35-36°C using a water-based heating pad (Gaymar Stryker T/Pump). Heart rate, body temperature and blood oxygenation level were continuously monitored for the duration of the experiments. All neural and EMG recordings were performed under ketamine anesthesia to avoid neural activity suppression. At the end of the experiment, the animals were deeply euthanized with a 120 mg/kg sodium pentobarbital intramuscular injection.

All studies were performed under the protocol approved by the UC San Diego Institution Animal Care & Use Committee (protocol S16020).

### Vertebrate Animal Subjects and Surgical Procedures for Chronic and Histological Studies

The electrode contacts were anchored on the vertebral bone, and guided from the implant target at the C4 spinal cord to the head underneath the skin. A 3-D printed implant made using ABS was fixed to the skull to make a skull-mount unit, which was held in place using dental cement. The thin film electrode was bonded to a custom acquisition PCB housed inside this skull mount unit.

We considered a stability threshold of 100 kΩ for a working electrode. Electrodes with impedances higher than this value would be unable to stimulate at high enough currents to cause neural activations. Our results showed that while the electrode impedances increased in a chronic setup, the electrodes remained functional in all 4 tested animals, with 15 out of the 16 contacts working at the time of explant, as shown in Fig. 5(D).

### Tissue Processing and Immunohistochemistry

#### Tissue processing

Subjects were deeply anesthetized and transcardially perfused with a 4% solution of paraformaldehyde, and the spinal cord was dissected out of the spinal column. Spinal cord dura was removed, and the spinal cord was cut in the transverse plane into 1cm blocks. The blocks of tissue were cut into 40-µm-thick transverse sections. Tissue sections were stored at –20 °C in tissue cryoprotectant solution (TCS; 25% glycerin (vol/vol).

#### Fluorescent immunolabelling

Transverse sections were pre-treated with 50% methanol for 20 min at 22–24 °C, washed in Tris-buffered saline (TBS) and blocked for 1 h in TBS containing 5% normal donkey serum and 0.25% Triton X-100. Sections were incubated in primary antibodies NeuN to label mature neurons (guinea pig, Millipore ABN90, 1:2,000,); c-FOS to label activated neurons (Rabbit, Santa cruz Biotech, sc-52, 1:200). Sections were washed with TBS and then incubated in Alexa-Fluor 488– or Alexa-Fluor 594– anti-donkey secondary antibodies (Thermo Fisher A11055, 1:500) for 1 h and DAPI (Sigma D9542, 0.5 µg/µl, to label nuclei) for 5 min. Sections were washed with TBS, mounted on slides, and coverslipped with Mowiol mounting medium.

#### Image acquisition

Epifluorescent images were captured using a Keyence BZ-X710 microscope and camera. For publication, images were processed uniformly for optimal brightness and contrast using Adobe Photoshop CS5 (Adobe Systems, San Jose, CA). Images intended for quantification (see below) were not modified in this fashion. Composites of multiple focal planes were constructed either in Adobe Photoshop CS5 (Wayne Rasband, National Institutes of Health; http://rsb.info.nih.gov/ij/).

### Quantification of c-FOS+ NeuN neurons in coronal sections

For the c-Fos analysis, c-FOS+, NeuN-labeled neurons were quantified in spinal segments C4 through C8. Counts represent the average of three sections per segment per rat. or ImageJ 1.48v (Wayne Rasband, National Institutes of Health; http://rsb.info.nih.gov/ij/).

Counts performed using A two-way ANOVA revealed a significant interaction between stimulation condition and spinal level (P < 0.0001). Analysis performed using PRISM software.

