## Supplementary Data for "Adaptive Charge Modulation Enables Focal, Selective Spinal Cord Stimulation"

### Supplementary Tables

| Muscle | p-value | Significance | Notes |
|---|---|---|---|
| Left Tibialis | 0.39 | - | |
| Left Bicep | 0.27 | - | |
| Left Vastus | 0.91 | - | |
| Left Semitendinosus | 0.19 | - | |
| Left Glutes | 0.52 | - | |
| Left External Oblique | 0.19 | - | |
| Right Gastro | 0.82 | - | |
| Right Tibialis | 0.55 | - | |
| Left Gastro | 0.62 | - | |
| Right Bicep | 0.03 | | No significant responses under stimulation |
| Right Vastus | 0.45 | - | |
| Right Semitendinosus | 0.99 | - | |
| Right Glutes | 0.60 | | |
| Right External Oblique | 0.97 | - | |

**Supplementary Table 1. Comparison of first response from offset vs no-offset stimulation.** The values shown here are for a two-sample t-test comparing the amplitude of the first response to stimulation for all activations of a given muscle. The first set is the amplitude in response to stimulation with no offset (both contacts stimulating at 1kHz). The second set is the amplitude in response to Adaptive Charge Modulation (the contacts stimulate at 1kHz and 967Hz). The data is for the July 11th, 2024 case.

| Muscle | p-value | Significance | Notes |
|---|---|---|---|
| Left Tibialis | 0.00029 | *** | |
| Left Bicep | 0.43 | - | No significant responses |
| Left Vastus | 0.22 | - | No significant responses |
| Left Semitendinosus | 0.0006 | *** | |
| Left Glutes | 0.079 | - | No significant responses |
| Left External Oblique | 0.91 | - | No significant responses |
| Right Gastro | 0.95 | - | No significant responses |
| Right Tibialis | 0.0017 | ** | |
| Left Gastro | 0.92 | - | No significant responses |
| Right Bicep | 0.0306 | * | |
| Right Vastus | 0.0000002 | *** | |
| Right Semitendinosus | 0.011 | * | |
| Right Gluteus Maximus | 0.7203 | - | No significant responses |
| Right External Oblique | 0.0047 | ** | |

**Supplementary Table 2. Comparison of subsequent responses, after the first response, from offset vs no-offset stimulation.** The values shown here are for a two-sample t-test comparing the amplitude of the non-first response to stimulation for all activations of a given muscle. The first set is the amplitude in response to stimulation with no offset (both contacts stimulating at 1kHz). The second set is the amplitude in response to Adaptive Charge Modulation (the contacts stimulate at 1kHz and 967Hz).

| Date | Weight | Type of Device | Experiment | Data Collected | Included in Manuscript |
|---|---|---|---|---|---|
| Oct 21, 2022 | 500g | 32 channel circumferential electrode | Pilot Experiment to Test Frequency Response | Yes | No; Pilot data superseded by subsequent experiments* |
| Nov 04, 2022 | 600g | SpineWrap electrode | Pilot Experiment for Validating Electrode Design | No | No |
| Nov 14, 2022 | 600g | SpineWrap electrode | Frequency Response of Lower Limb EMG Responses | Yes | No; Preliminary data superseded by subsequent experiments* |
| Jan 24, 2023 | 550g | ECAP Electrode v01 | Parameter Space Exploration of EMG Responses | No | No |
| Feb 01, 2023 | 550g | ECAP Electrode v01 | Parameter Space Exploration of EMG Responses | Yes | No; Preliminary data superseded by subsequent experiments* |
| Feb 14, 2023 | 600g | ECAP Electrode v01 | Frequency Response of Lower Limb EMG: Sinusoidal Waves | No | No |
| Feb 22, 2023 | 600g | ECAP Electrode v01 | Frequency Response of Lower Limb EMG: Sinusoidal Waves | Yes | Yes; Included in Supplementary Data |
| Apr 19, 2023 | 800g | ECAP Electrode v02 | Frequency Response of Lower Limb EMG + ECAP | Yes | No; Preliminary data superseded |

| | | | | | |
|---|---|---|---|---|---|
| | | | | | by subsequent experiments* |
| May 3rd, 2023 | 850g | ECAP Electrode v02 | Frequency Response of Lower Limb EMG + ECAP | No | No |
| May 16th, 2023 | 800g | ECAP Electrode v02 | Frequency Response of Lower Limb EMG + ECAP | Yes | No; Preliminary data superseded by subsequent experiments* |
| May 18th, 2023 | 900g | ECAP Electrode v02 | Frequency Response of Lower Limb EMG + ECAP | Yes | No; Preliminary data superseded by subsequent experiments* |
| Jan 12th, 2024 | 950g | ECAP Electrode v03 + 1024-Channel ECoG Grid v01 | Frequency Response of Lower Limb EMG + ECAP + Cortical Responses | Yes | Yes |
| Mar 14th. 2024 | 900g | ECAP Electrode v03+ 1024-Channel ECoG Grid v01 | Frequency Response of Lower Limb EMG + ECAP + Cortical Responses | Yes | Yes |
| Apr 30th, 2024 | 1100g | Concentric Ground Electrode + 1024-Channel ECoG Grid v01 | Frequency Response of Lower Limb EMG + ECAP + Cortical Responses | Yes | Yes |
| Jun 21st, 2024 | 950g | Concentric Ground Electrode + 1024-Channel ECoG Grid v02 | Sensory Mapping of Target EMG Muscles | Yes | Yes |
| Jul 11th, 2024 | 1050g | Concentric Ground Electrode + 1024-Channel ECoG Grid v02 | Adaptive Charge Modulation Experiment 1 | Yes | Yes |
| Dec 10th, 2024 | 900g | Concentric Ground Electrode + 2 x | Adaptive Charge Modulation Experiment 2 | Yes | Yes |

| | | |
|---|---|---|
| | | 1024-Channel ECoG Grid v02 |

**Supplementary Table 3. Summary of Rat Experiments for Demonstration of ACM.** All the datasets used in the experiment will be made available. *Dataset included in Supplementary Information.

### Supplementary Figures

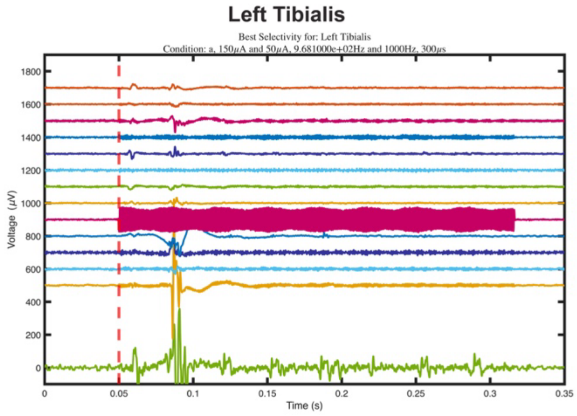
Left Tibialis
Best Selectivity for: Left Tibialis
Condition: a, 150μA and 50μA, 9.681000e+02Hz and 1000Hz, 300μs
Voltage (μV)
Time (s)

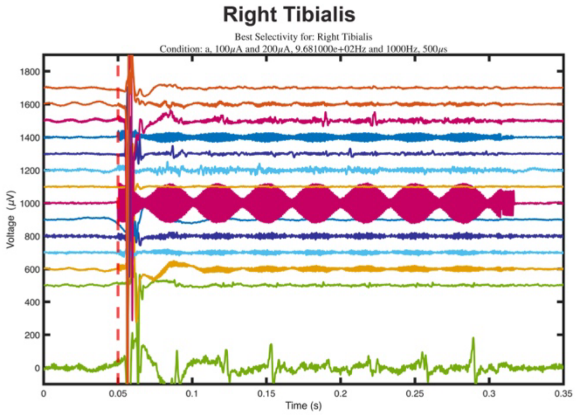
Right Tibialis
Best Selectivity for: Right Tibialis
Condition: a, 100μA and 200μA, 9.681000e+02Hz and 1000Hz, 500μs
Voltage (μV)
Time (s)

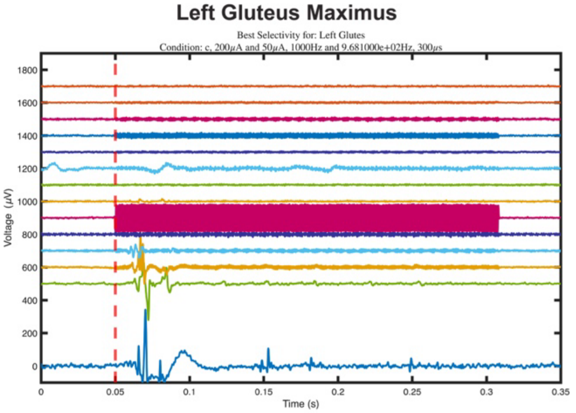
Left Gluteus Maximus
Best Selectivity for: Left Glutes
Condition: c, 200μA and 50μA, 1000Hz and 9.681000e+02Hz, 300μs
Voltage (μV)
Time (s)

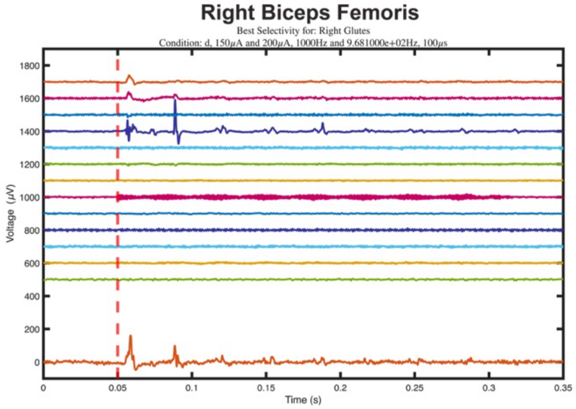
Right Biceps Femoris
Best Selectivity for: Right Glutes
Condition: d, 150μA and 200μA, 1000Hz and 9.681000e+02Hz, 100μs
Voltage (μV)
Time (s)

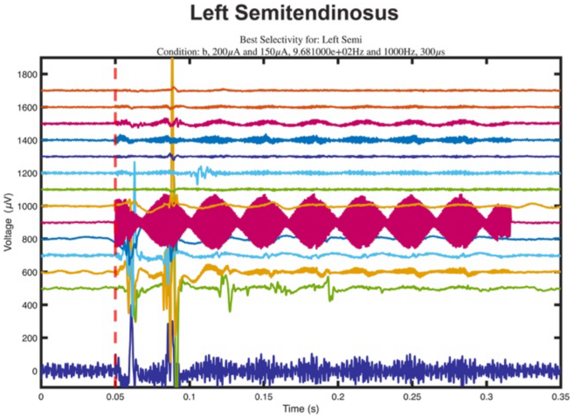
Left Semitendinosus
Best Selectivity for: Left Semi
Condition: b, 200μA and 150μA, 9.681000e+02Hz and 1000Hz, 300μs
Voltage (μV)
Time (s)

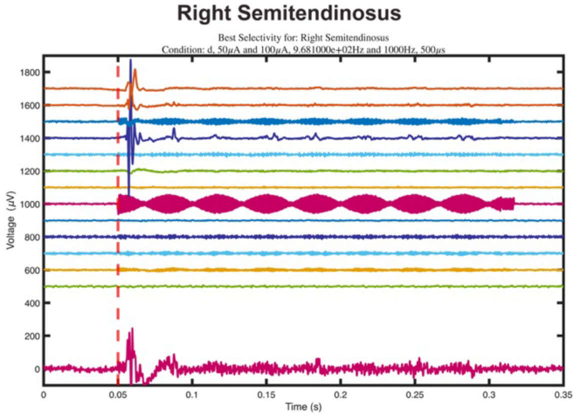
Right Semitendinosus
Best Selectivity for: Right Semitendinosus
Condition: d, 50μA and 100μA, 9.681000e+02Hz and 1000Hz, 500μs
Voltage (μV)
Time (s)

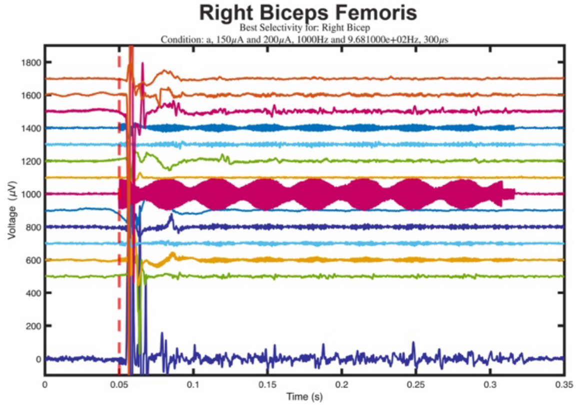
Right Biceps Femoris
Best Selectivity for: Right Bicep
Condition: a, 150μA and 200μA, 1000Hz and 9.681000e+02Hz, 300μs
Voltage (μV)
Time (s)

**Supplementary Figure 1: Maximum Selectivity of Responses Plots:** We achieved SI =1 for these 7 muscles within one experiment, demonstrating the strength of the Adaptive Charge Modulation approach.

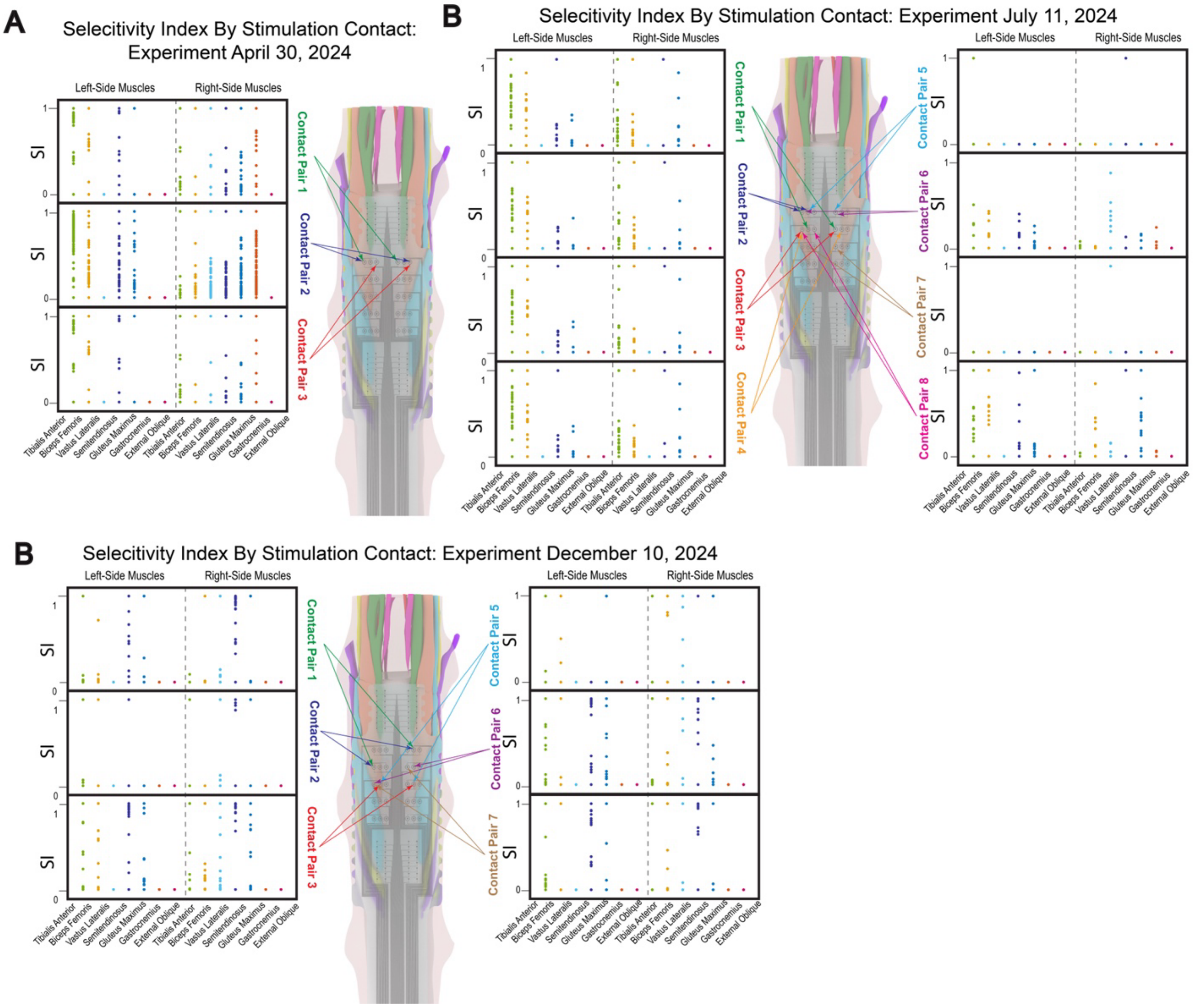

**Supplementary Figure 2: Selectivity Index Achieved in Each Experiment.** (A) Selectivity Index from 3 contact pairs stimulated on the April 30, 2024 experiment. (B) Selectivity Index from 8 contact pairs stimulated on the July 11, 2024 experiment. The left 4 panels are the same as shown in Fig. 3(B) of the main text. (C) Selectivity Index from 3 contact pairs stimulated on the Dec 10, 2024 experiment. Each plot shows the selectivity indices achieved across multiple stimulation conditions for the same contact. This is an overview of the achieved selectivity from all stimulation conditions- the selectivity from a single condition is represented on radial plots in other figures.

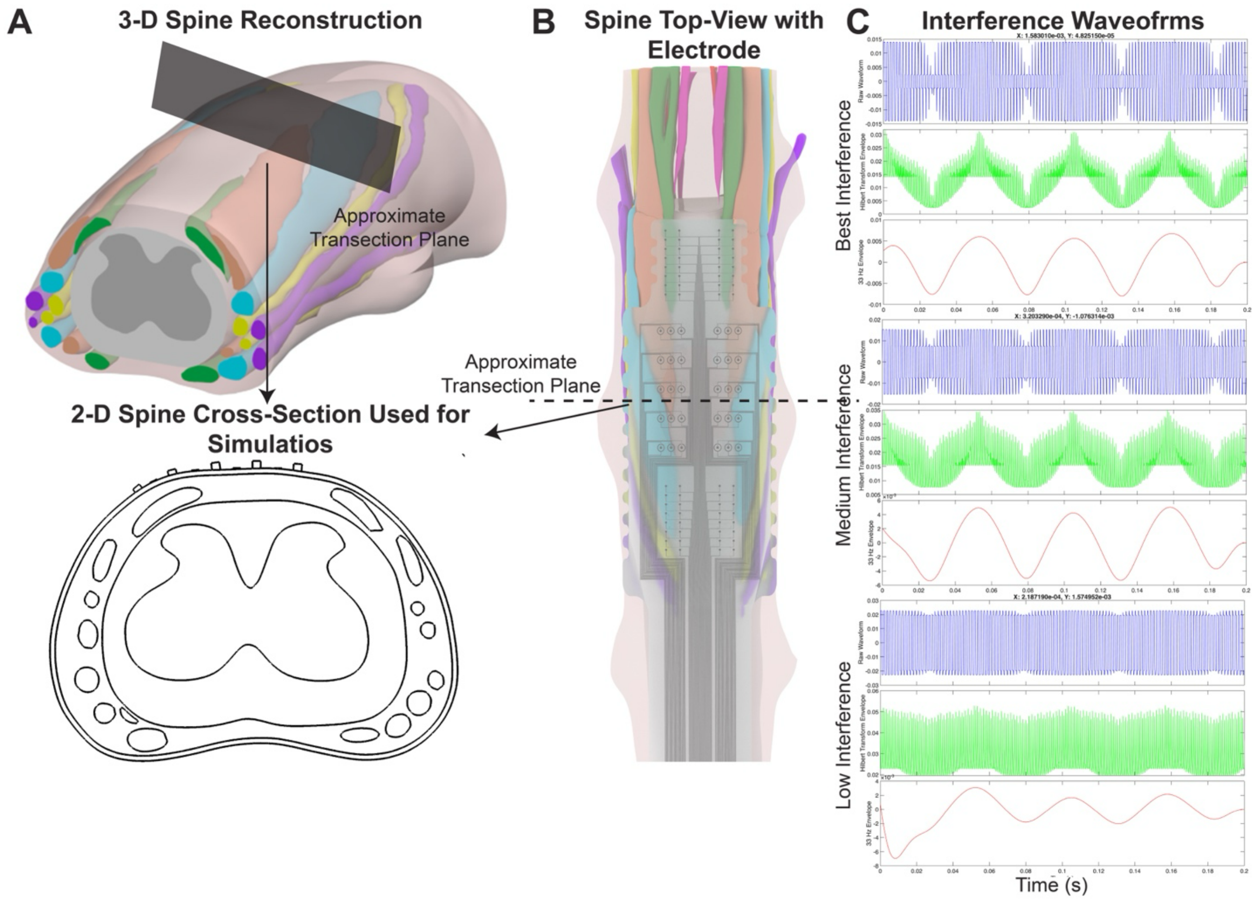

**Supplementary Figure 3: Simulation Setup for COMSOL Simulations.** (A) The COMSOL 3-D spinal reconstruction used for simulations, and the approximate transection plane used for the 2-D cross section showed in simulations. (B) Top-view of the concentric ground electrode placed on the spine, approximated at the T13-L1 junction. (C) Interference ratio represented in the simulations. The first row in each plot is the raw waveform, the second row is the Hilbert Transform of the raw voltage, and the third plot is the 10-50Hz band-passed wave approximating the modulation.

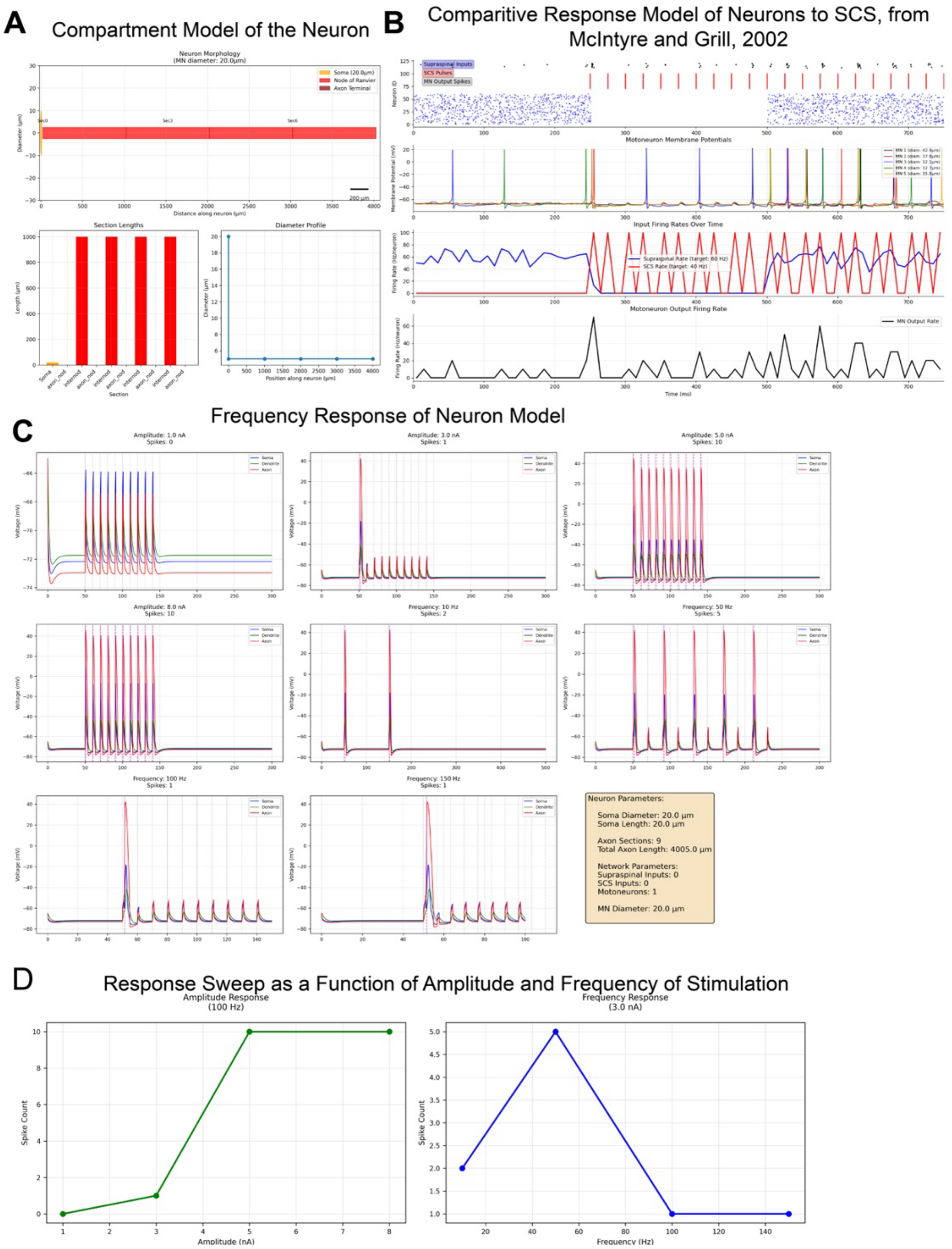

**Supplementary Figure 4: NEURON model used for simulation of responses shown in Fig. 1(D).** (A) The compartment model and size of each neuron component. (B) Recreated responses to Spinal Cord Stimulation (SCS), using Hodgkin-Huxley simulation parameters reported in McIntyre and Grill, 2002. (C) The time-domain neuron simulation response as a function of frequency. (D) The relative activation as a function of amplitude and frequency of stimulation.

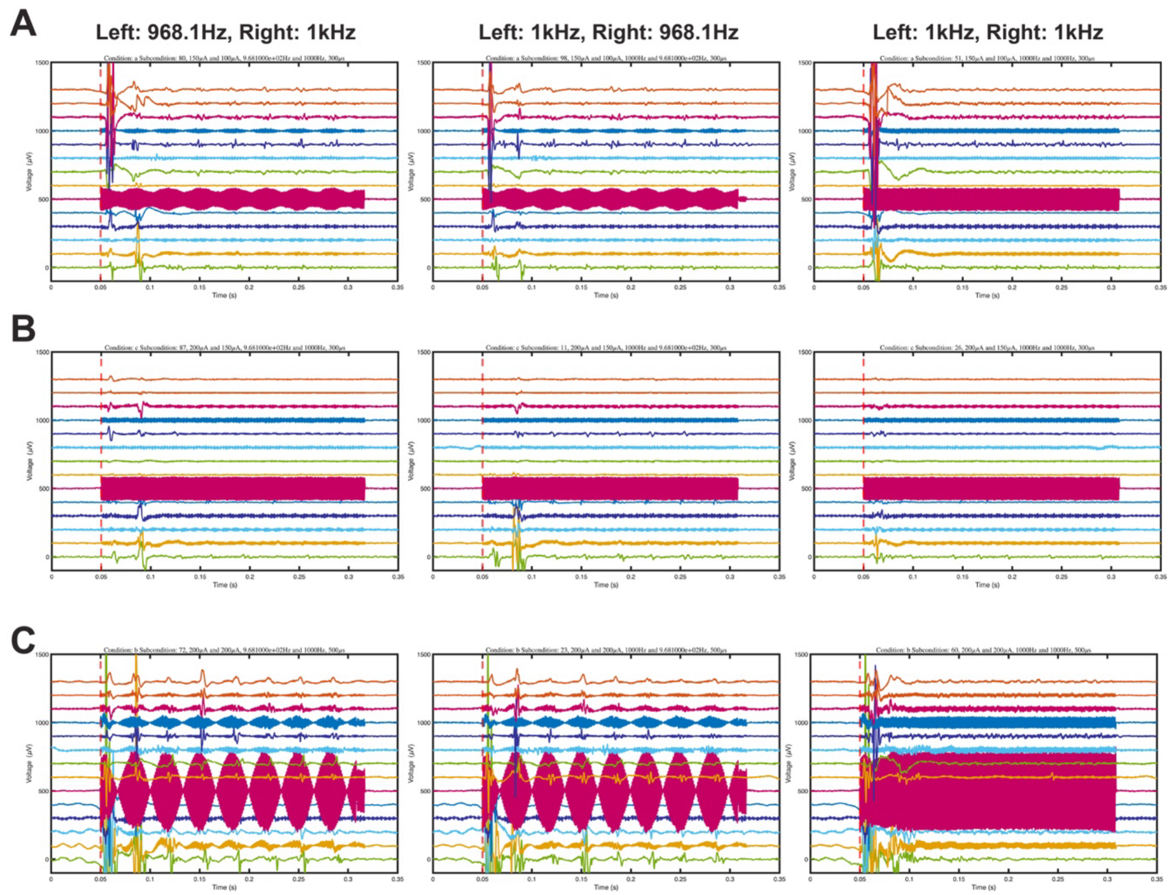

**Supplementary Figure 5: Demonstration of Adaptive Charge Modulation.** For each stimulation parameter combination, the contacts were stimulated with offset (968.1Hz and 1kHz, and 1kHz and 968.1Hz), and without offset (1kHz and 1kHz). This figure shows the responses with and without frequency offset, demonstrating the responses arise due to offset in frequency alone (all other parameters are kept constant). (A) Contact 1: 150µA, Contact 2: 100µA, Pulse Width: 300µs, (B) Contact 1: 200µA, Contact 2: 150µA, Pulse Width: 300µs, (C) Contact 1: 200µA, Contact 2: 200µA, Pulse Width: 500µs.

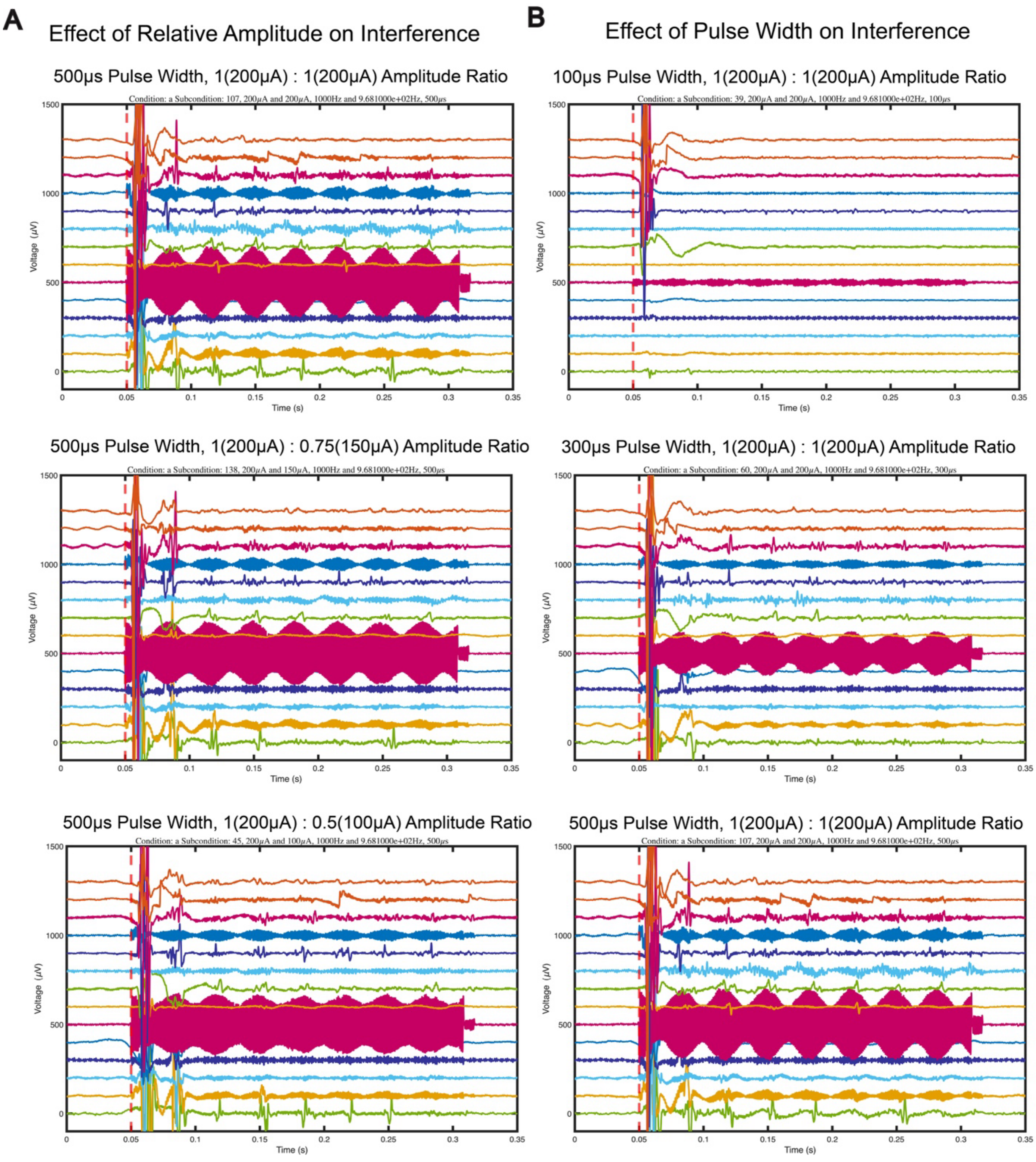

**Supplementary Figure 6: EMG Plots Corresponding to the Selectivity Index Plots shown in Figure 2(B) and 2(C).**

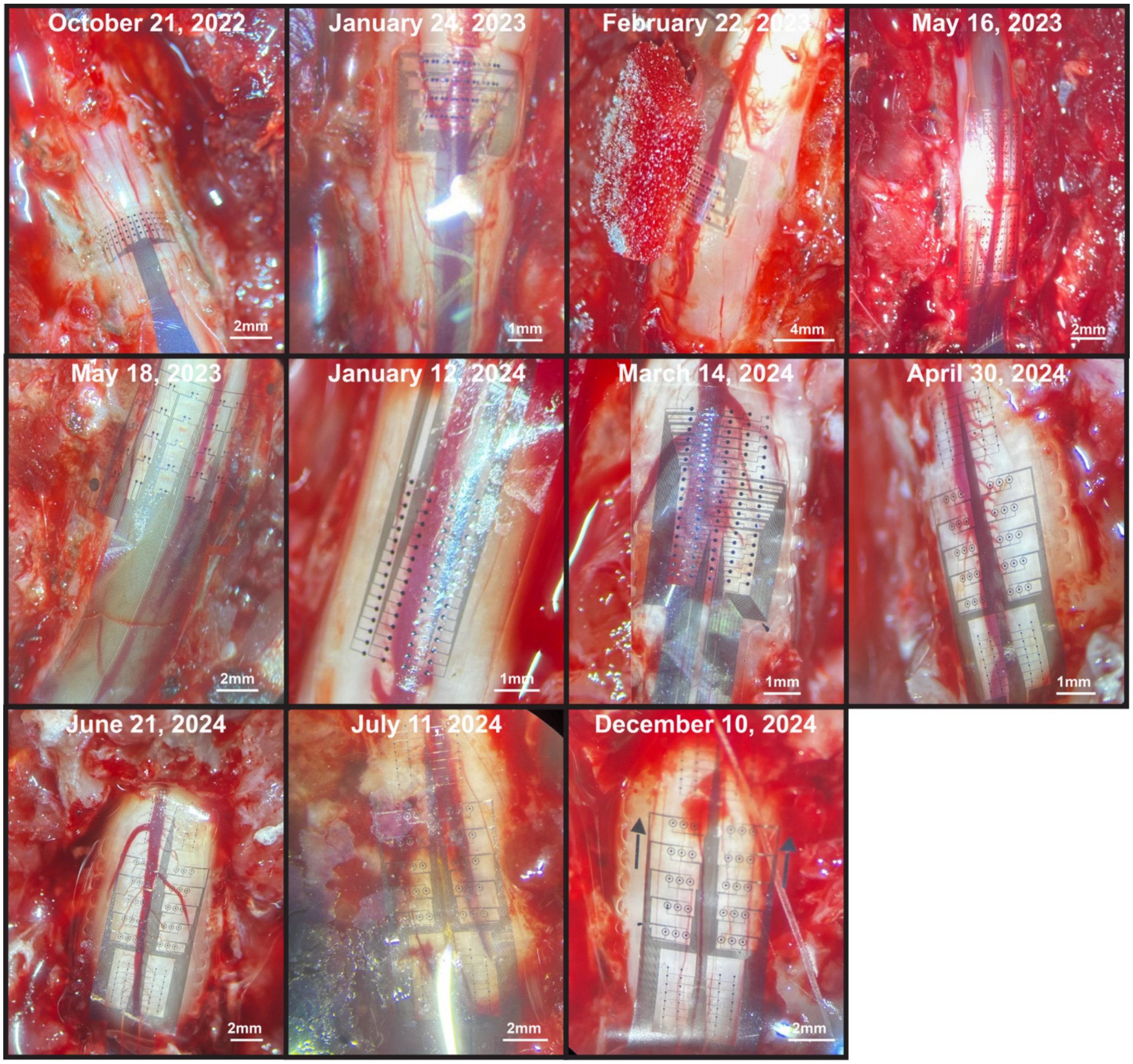

**Supplementary Figure 7. Electrode Placement on the Spine Across Experiments**. The top of each figure represents the rostral direction.

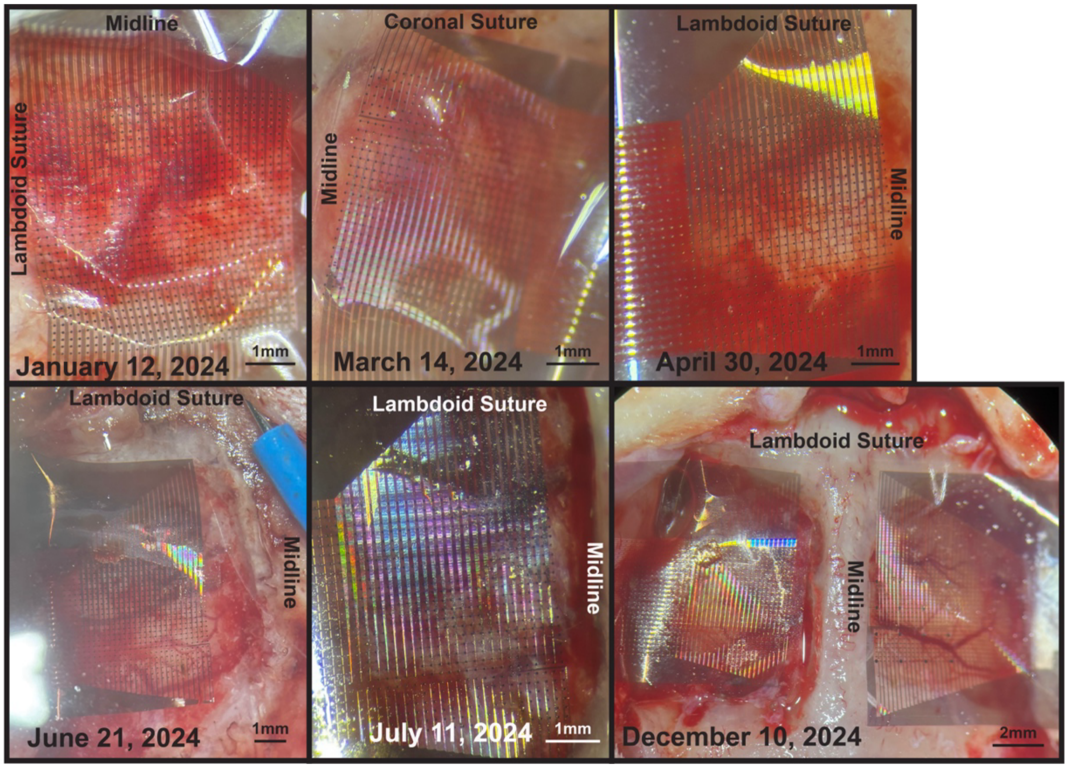

**Supplementary Figure 8. Electrode Placement on the Brain Across Experiments**.

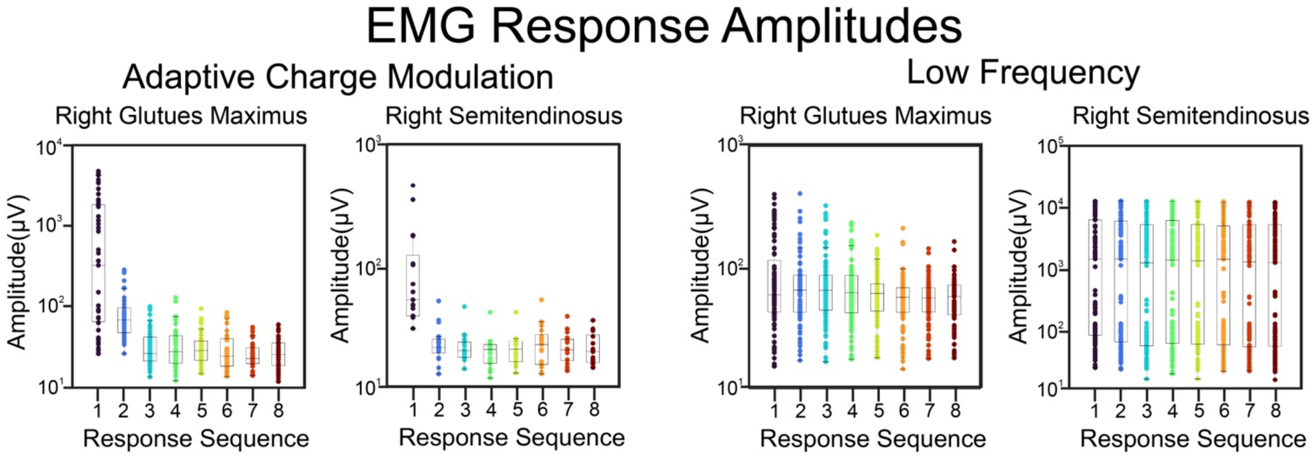

**Supplementary Figure 9. EMG Response Amplitude Profiles.** The response amplitudes, as a function of the response sequence number. The initial response typically has the highest amplitude, with decaying amplitude for subsequent responses.

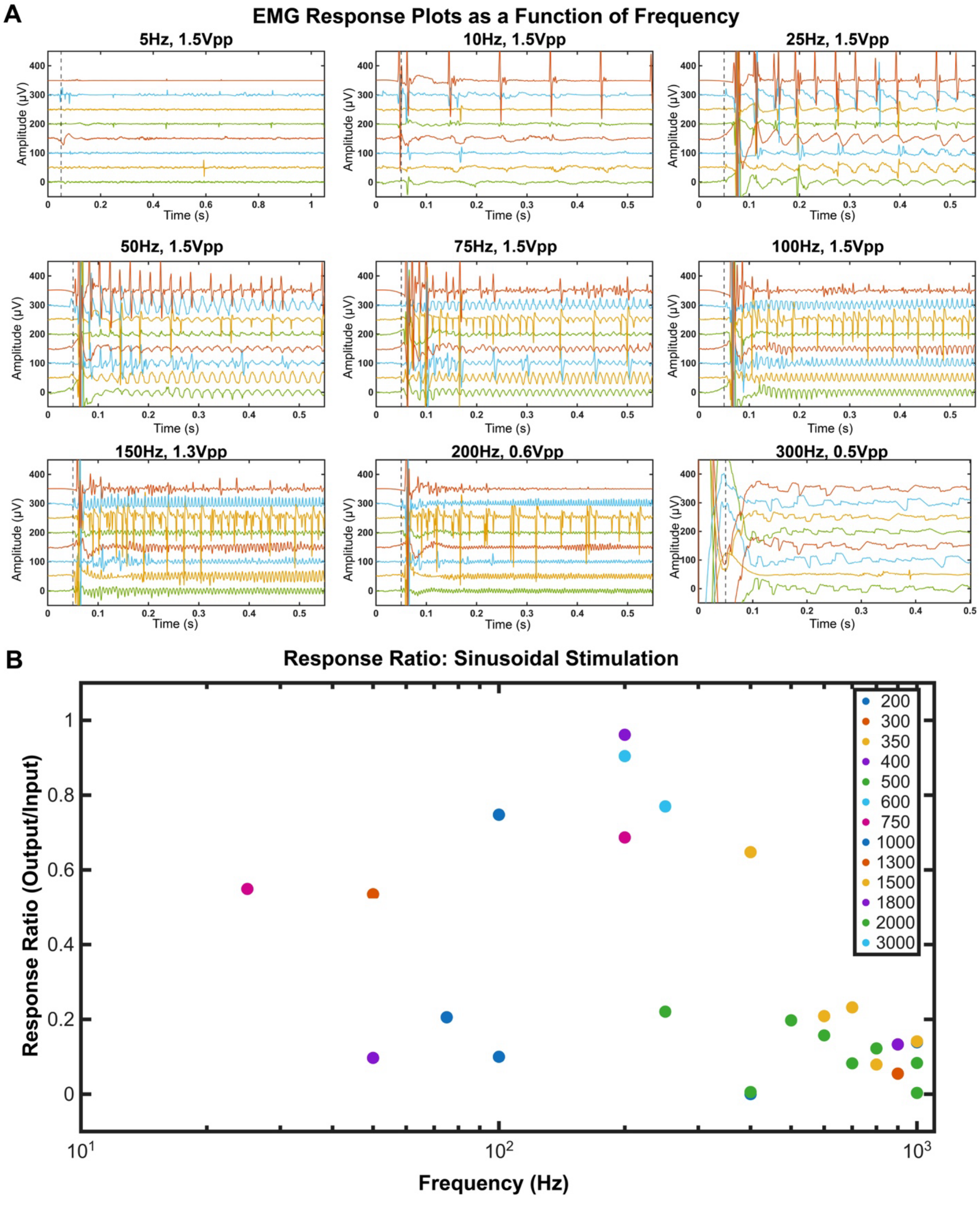

**Supplementary Figure 10: Sinusoidal Stimulation Responses as a Function of Frequency of Sinusoidal Stimulation Wave.** (A) The individual panels show the measured EMG response for 8 measured muscles, to continuous, voltage-clamped sinusoidal stimulation. The 8 muscles measured are, from top to bottom in each panel: Left Tibialis Anterior, Left Bicep Femoris, Left Vastus Lateralis, Left Gastrocnemius, Right Tibialis Anterior, Right

Bicep Femoris, Right Vastus Lateralis, Right Gastrocnemius. (B) The response ratio for stimulation to the sinusoidal wave, showing decay in response with higher frequency.

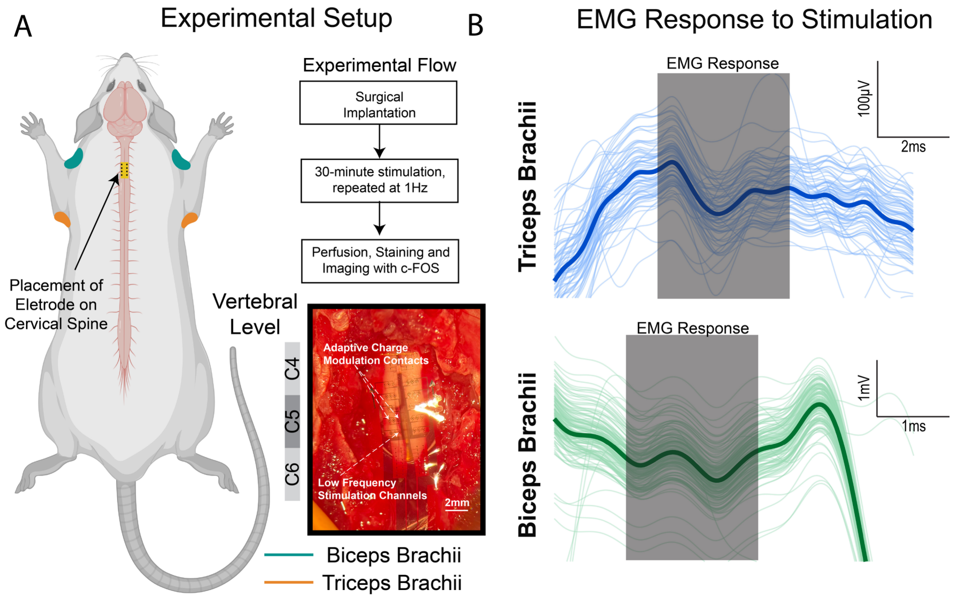

**Supplementary Figure 11. Acute histological imaging experiments setup.** (A) Experimental setup for the acute experiments, showing the placement of the electrode on the spine, and the target upper limb muscles. (B) The actual measured EMG response to stimulation.

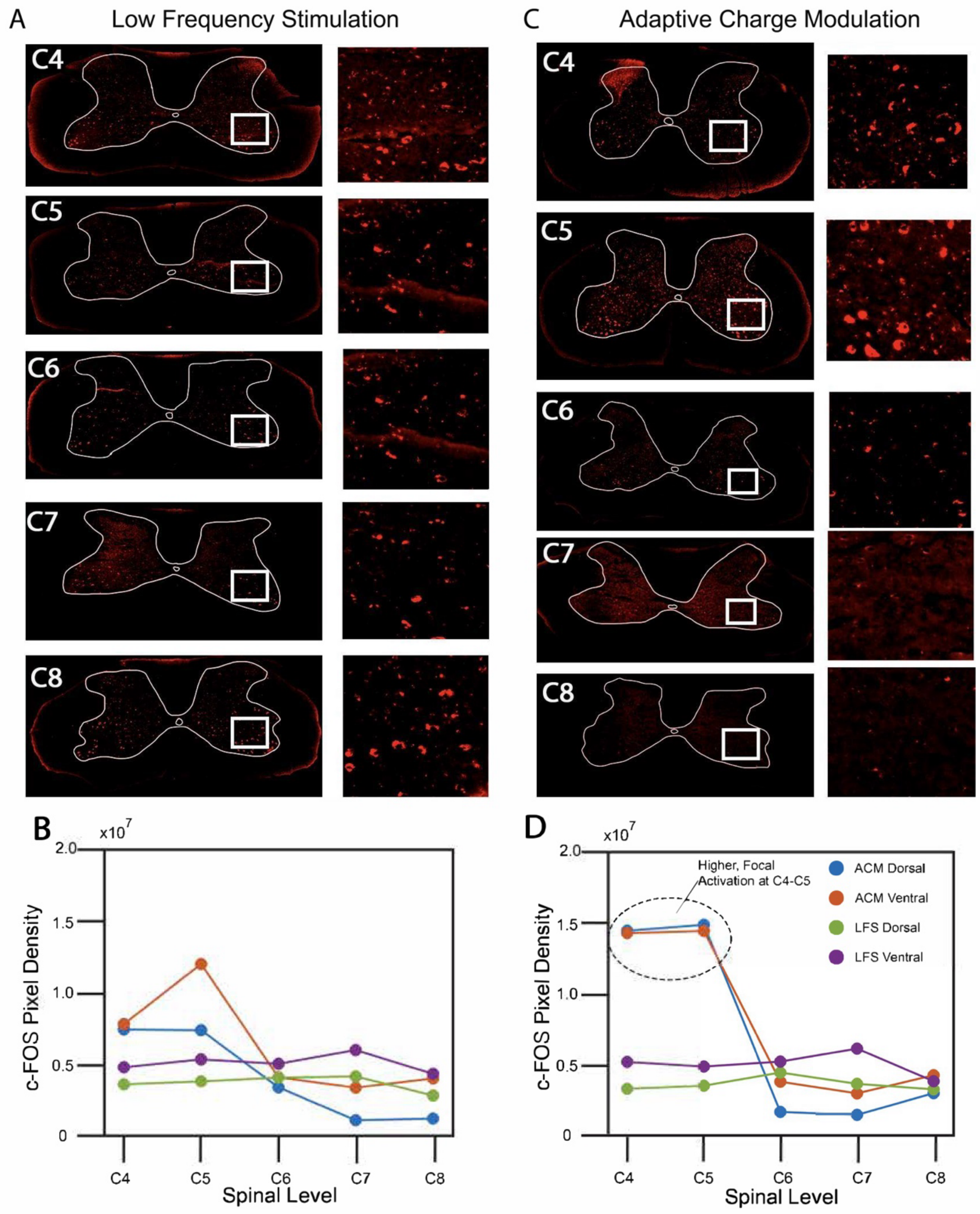

**Supplementary Figure 12. Acute histological imaging results.** The c-FOS-stained images imaged after sacrificing the animal, for (A) low frequency stimulation and (C) adaptive charge modulation. The measured cFOS expression intensity in the left and right dorsal and ventral columns of the spine, for (B) low frequency stimulation and (D) adaptive charge modulation.

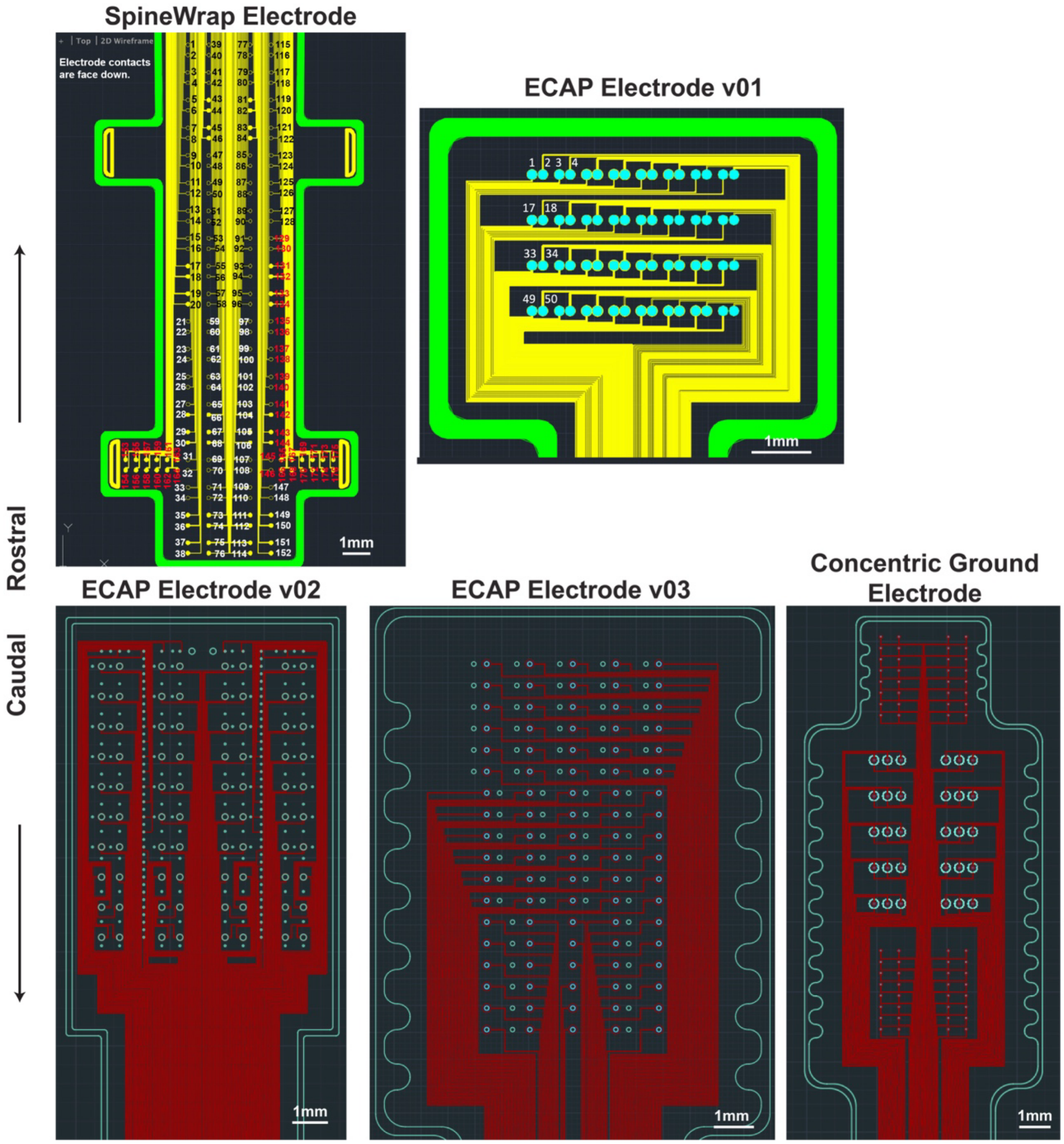

**Supplementary Figure 13. Spinal Electrode Designs used across experiments, demonstrating the evolution of the design across experiments**. The design names correspond to the "Type of Device" Column in Supplementary Table 3.

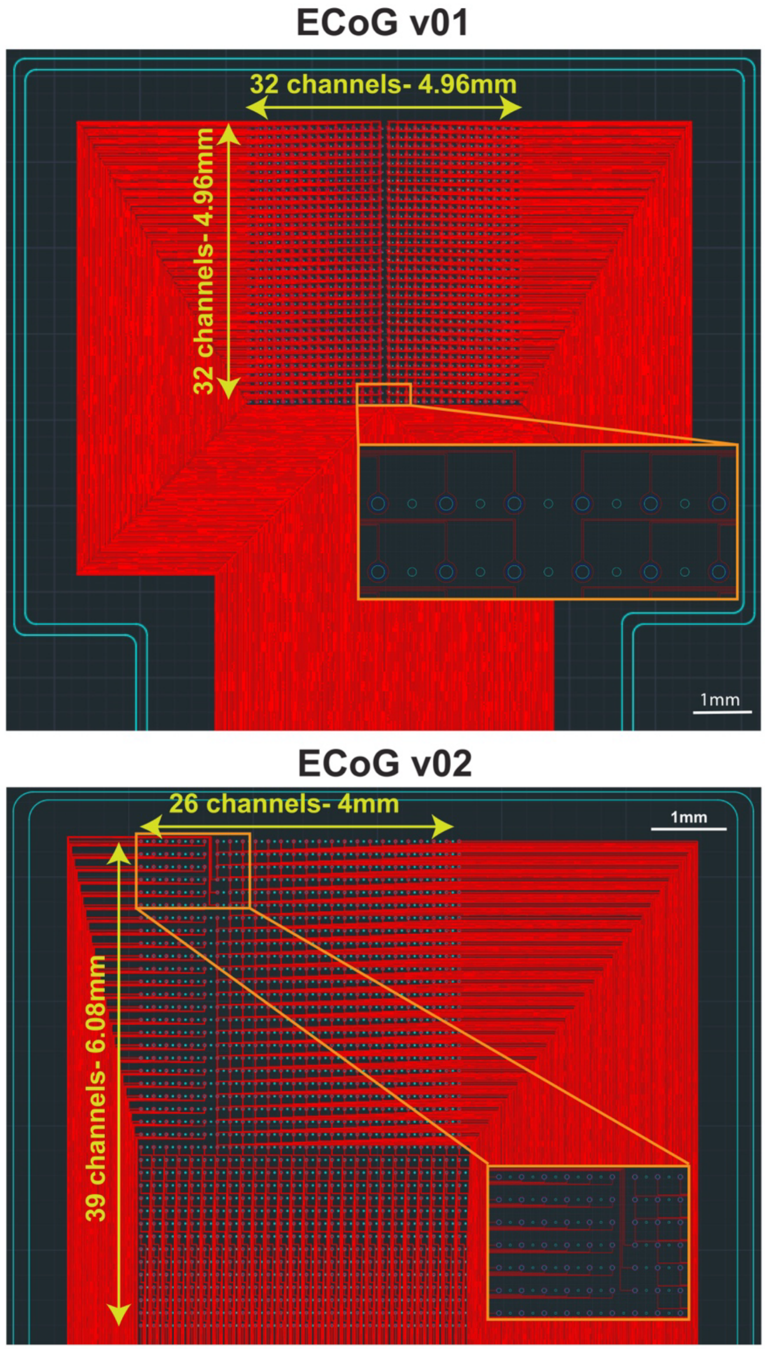

**Supplementary Figure 14. Cortical Electrode Designs used across experiments, demonstrating the evolution of the design across experiments**. The design names correspond to the "Type of Device" Column in Supplementary Table 3.
